# Wildfires during fall migration affect bird movements, avian health, and migration timing in western North America: Insights from bird banding and feather isotopes

**DOI:** 10.64898/2026.01.27.701926

**Authors:** Kyle D. Kittelberger, Colby J. Tanner, Gabriel J. Bowen, Llewellyn W. Stringer, Çağan Hakkı Şekercioğlu

## Abstract

With the peak of the wildfire season in North America typically occurring during the core part of fall bird migration, migratory birds are likely to be increasingly impacted by worsening wildfire seasons during their southerly movements. In this study, we combined two approaches to assess the impact of wildfires on migratory birds captured in southern Utah. First, we used five years of bird banding data to assess how bird movement and physical condition patterns are impacted. Second, we used stable hydrogen isotopes from collected feathers to identify the geographic origin of the individuals of several migratory species in order to better understand how fires may be influencing migration. We found that when there were more wildfires active in western North America, there were more captures of birds at our banding station, likely a result of birds shifting their movements to avoid areas of fire and smoke, and that birds had worse body condition and overall health as shown from lower body masses. We also found that during periods when wildfires are especially active and severe, wildfire activity has the most significant influence on bird movements and health compared to other prominent environmental variables. Our isotopic results provide evidence that two species’ migration patterns varied across the year in both migration timing and likely summer origin, and show how fire activity could have affected the migratory window for some species. More broadly, our isotope data improves our knowledge of the likely geographic origins of migratory birds in this particular flyway, adds to the literature on feather-based hydrogen isotope analyses, provides some of the first published isotope data for two species, and streamlines a workflow that can aid future researchers working with feather hydrogen isotope data.

## 1. INTRODUCTION

Billions of birds annually migrate between their breeding and non-breeding grounds around the world (Dokter et al. 2018), utilizing stopover sites along their migratory routes to aid in the successful completion of these long-distance journeys (Mehlman et al. 2005, Blount et al. 2021, Roques et al. 2022). However, recent studies have revealed significant declines in bird populations, particularly of migratory species, on continental scales over the past few decades (Sanderson et al. 2006, Vickery et al. 2014, Horns and Şekercioğlu 2018, Rosenberg et al. 2019, Burns et al. 2021). Climate change is one of the major threats to migratory birds (Wormworth and Şekercioǧlu 2011, Abolafya et al. 2013, Kittelberger et al. 2022), with a majority of forest birds in the western, boreal zones of North America particularly vulnerable to rising temperatures (Bateman et al. 2020) and changes in precipitation regimes (Illán et al. 2014). In western North America, climatic warming has been contributing to extreme persistent droughts and more severe wildfire seasons (Abatzoglou and Williams 2016, Williams et al. 2020, Kittelberger et al. 2022, Zhai et al. 2022), both of which can function as key stressors for birds (Saracco et al. 2018, Cady et al. 2019, Kittelberger et al. 2022, Irannezhad et al. 2022, Stanek et al. 2025).

With the peak of the wildfire season typically occurring during the core part of fall bird migration, migratory birds in particular are likely to be increasingly impacted by worsening wildfire seasons across western North America during their southerly movements (Kittelberger et al. 2022, Overton et al. 2022, Nihei et al. 2024). In fact, during the fall of 2020, there was a series of widespread mass die-off events of thousands of migratory birds in the western United States that coincided with the second most severe wildfire season on record for the region (Kittelberger et al. 2022, Irannezhad et al. 2022, NIFC 2025). One study found that some birds can significantly alter their migratory movements and be forced into other habitats in order to avoid flying through extensive wildfire smoke (Overton et al. 2022). Additionally, analyses of bird banding data from a key migratory stopover site in southeastern Utah revealed that during fall 2020, more bird captures were correlated with more acres burned by wildfires for the day birds were captured (Kittelberger et al. 2022). Furthermore, a reduction in body mass of captured birds in fall 2020 was correlated with more acres burnt one week prior to capture (Kittelberger et al. 2022). Birds are known to detect and move ahead of oncoming natural disasters (Chilson et al. 2012, Streby et al. 2015, Heckscher 2018), so it is possible that extreme fires, such as those in 2020, may force birds to prematurely begin their migration before they have stored all of the necessary fat reserves for such a journey (Pennisi 2020, Kittelberger et al. 2022, Irannezhad et al. 2022). Furthermore, body mass reductions have been linked to exposure to wildfire smoke in southern California (Nihei et al. 2024). Burnt stopover sites can cause birds to continue flying in search of habitat and food or skip stopovers altogether and fly as far south as possible (Frost 2018), but wildfire smoke can also cause birds to increase their number of stopovers (Overton et al. 2022). However, there is still relatively limited research examining how wildfires affect birds during migration, in part due to the challenge of spatially and temporally connecting wildfires with the responses of birds that migrate (Kittelberger et al. 2022, Overton et al. 2022).

Knowing where birds are migrating from can help better understand the impacts of fires and wildfire smoke on bird migration. Although the large majority of Neotropical songbirds have body masses too small to allow tracking through conventional GPS satellite transmitters, isotopic analyses of keratin in feather samples can provide an effective proxy (Hobson and Wassenaar 1997, Kelly and Finch 1998, Hobson et al. 2014). Isotopes can help link actively migrating birds with their breeding ground locations (Kelly and Finch 1998, Hobson et al. 2014, 2015, Brattström et al. 2018, Wommack et al. 2020), indicating where birds migrated from and reveal the general route they took. Specifically, the deuterium (heavy hydrogen) isotope has been widely used as a geographic marker because there are well-known latitudinal gradients in deuterium to hydrogen ratios (δ^2^H) in rainfall across the landscape (Bowen et al. 2005). This enables an estimation of where a bird has been via the isotopic analyses of collected feathers (Bowen et al. 2005, Norris et al. 2006, Bowen 2010, Hobson et al. 2015), as the ratio of deuterium isotopes in the sample will reflect the isotopic signature of precipitation where the feather was formed. This can therefore allow us to determine where a bird likely bred during the summer prior to its fall migration. We could then use this knowledge to assess whether the migration of individual birds from specific geographic areas differs across years and, if so, whether this could be due to the effects of wildfires.

In this study, we examined the impacts of wildfires on birds during fall migration in western North America across a multi-year period using two main approaches. First, we utilized bird banding data from southern Utah to assess how bird movement and condition patterns, assessed via the number of daily bird captures and an individual’s body mass, are impacted in relation to the extent of average wildfire acres burned in western North America across years. We used data from a five-year period that spanned from 2019 to 2023 in order to include the years before and after the severe wildfire season and notable avian mass mortality events of 2020 (Kittelberger et al. 2022). Additionally, to further parse out the effects of wildfires in our study from that of other important environmental factors, we also included precipitation, temperature, and drought in our analyses. We predicted that across years, we would see more bird captures and more under-weight birds when there were more wildfire acres burning.

For our second approach, we sought to identify the geographic origin of the individuals of several migratory species that seemed to have been impacted physically by wildfires, in order to better understand how fires may be influencing fall migration. To do this, we conducted stable hydrogen isotope analyses on feathers collected in southern Utah to examine if birds may be shifting departure dates from certain areas over time due to wildfires. We predicted that birds from similar geographic areas may initiate their migration earlier in years that had a higher risk of wildfires. To our knowledge, this study represents the first assessment of avian responses to wildfires using stable hydrogen isotopes of feathers.

## 2. METHODS

### 2.1 Study site

Fieldwork for this research took place at the Bonderman Field Station at Rio Mesa (38.799 °N, 109.205 °W) in Grand County, Utah, United States, a restricted-access research site that is property of the University of Utah. This station is situated in the American Interior West, near the boundary of the Western/Pacific and Central migratory bird flyways of North America (La Sorte et al. 2014, Rosenberg et al. 2019, Kittelberger et al. 2022, 2025, Allocca et al. 2025). At an elevation of 1280 m asl, the research station is situated along the Dolores River and within the arid canyonlands region of the Colorado Plateau (Kittelberger 2021). Mist-nets for birds are placed within the strip of riparian vegetation that borders the river on one side and more open field-type or sagebrush habitats on the other side (Kittelberger et al. 2022, 2025, Allocca et al. 2025). The vegetation here consists of a mixture of native and non-native trees and shrubs, with the predominant species being tamarisk (*Tamarix chinensis*), skunkbush (*Rhus trilobata*), New Mexico privet (*Forestiera pubescens*), Fremont cottonwood (*Populus fremontii*), coyote willow (*Salix exigua*), and Goodding’s willow (*Salix gooddingii*).

### 2.2 Bird banding

Bird banding has taken place annually each fall at Rio Mesa since 2011, with the seasons typically spanning mid-to-late August through early November in order to encompass the entirety of fall land bird migration in eastern Utah (Kittelberger et al. 2022). For more information on the mist-net methodology employed at Rio Mesa, see Kittelberger et al. (2022, 2025).

Once extracted from nets, birds were first identified to species before an aluminum leg band (Bird Banding Laboratory, USGS) was placed on the legs of newly captured birds. For recaptured individuals previously banded, the band numbers were documented. The age and sex of individuals were then determined if possible (Pyle 1997, 2022). Physiological and morphological data were then recorded by experienced bird banders, including fat score, body mass, and body condition. For body mass, birds were weighed on digital scales to the nearest 0.1 grams. Fat score is measured on a scale of 0 to 7 and reflects the amount of fat deposited on the body of a bird. Body condition, recorded since 2021, is scored on a scale of 1 to 3 and reflects the degree of potential emaciation based on the shape of the breast muscles (see Kittelberger et al. (2022) for detailed methodology). Finally, one or two tail feathers (rectrices) are carefully collected from a bird before being placed inside an envelope with associated metadata noted. These feathers were first kept in a freezer at the banding station and then stored in a 4°C cold room in the Şekercioğlu Laboratory at the University of Utah following the completion of each banding season.

### 2.3 Dataset

#### 2.3.1 Bird banding data

We removed the records of any birds that were either not identified to the species level or were noted as of hybrid origin. As we chose to follow taxonomic classifications based on the eBird/Clements Checklist of Birds of the World (Clements et al. 2025), we lumped the following groups at the species level: Red-shafted and intergrade forms of Northern Flicker (*Colaptes auratus*); Gambel’s and Mountain subspecies of White-crowned Sparrow (*Zonotrichia leucophrys*); Gray-headed, Oregon, Pink-sided, Cassiar, and Slate-colored forms of Dark-eyed Junco (*Junco hyemalis*); and Audubon’s and Myrtle subspecies of Yellow-rumped Warbler (*Setophaga coronata*).

Finally, as we were focusing on the effects of wildfires in this study, particularly in how the impacts of the 2020 wildfire season on bird banding in the fall of that year compared with the impacts in other years. Because we needed to develop a multi-year daily wildfire dataset that could be used in analyses with migratory bird data (Kittelberger et al. 2022), we filtered our banding data to the fire year period spanning 2019 to 2023. We then standardized our data across years by restricting the fall seasons to banding days between August 31 and November 3 (Kittelberger et al. 2022), as this was the season length during fall 2020. Additionally, we removed the data of one nocturnal owl, as our focus for this study was on diurnal species. This resulted in a five-year fall Rio Mesa banding dataset consisting of 85 species comprising 3396 captures (Appendix A). None of the birds in our dataset were recently fledged juveniles (e.g., birds undergoing juvenile flight feather molt).

We then created separate datasets for overall number of captures, body mass, and body condition. For mass, we first filtered out any birds that either lacked recorded body masses and/or had no fat scores recorded (n=300). We checked the data for individual birds that were more than five standard deviations from each species’ mean (Youngflesh et al. 2022) and corrected any mistakes; we removed any values from our analyses that we were not able to correct (n=2). This resulted in a body mass dataset consisting of 3094 captures. Likewise, for body condition, we filtered out any birds that lacked this value. Since body condition was only collected beginning in 2021, our dataset comprised of 3 years of data, resulting in a dataset consisting of 1532 captures.

#### 2.3.2 Wildfire Data

To develop a dataset of estimated daily wildfire acres burned, we followed the methods established by Kittelberger et al. (2022). We retrieved spatial shapefile data on wildfires in the western United States from the open source platform National Interagency Fire Center (2019), and from the Canadian Forest Service (2021) for data on fires in western Canada. We then used ArcGIS Pro (ESRI 2021) to select fires in our five-year study period from the same western North American study region examined by Kittelberger et al. (2022). We then chose fires that were active at any point during the banding season as well as up to two weeks before the start of the season, in order to account for the hyperphagia period in which birds are depositing fat in preparation for migration (Driedzic et al. 1993, Morton and Pereyra 1994, Bairlein 2002, La Sorte et al. 2018, Kittelberger et al. 2022). For more details on the rationale for selecting our spatial study region and temporal period for data, see Kittelberger et al. (2022).

Next, after selecting fires of size classes E, F, or G (NWCG 2021) in order to focus on those that were likely to have had an important impact on habitat and wildlife (Kittelberger et al. 2022), we used the name or identifier for each fire to research publicly available information on the duration of these incidents. When both containment and control dates were differentiated for a fire (and often they are the same when control dates are also noted), and to try to standardize duration across fires and years, we chose to utilize containment dates in order to focus on the main period in which a fire was burning and spreading (since after containment, the size of a fire should not significantly change). When no information could be found about the end date of a fire, we used our best judgement to make a conservative evaluation (e.g., the last reported update on a fire). We also updated the dataset for the fires in 2020 compared to what was in Kittelberger et al. (2022), including some fires that were not originally part of the national dataset for that year. We used this process to double-check that we were only including fires that were active between August 23 and November 3. Additionally, there were some fires from 2021 and 2022 from British Columbia for which we could not find complete information on their duration, so we removed these (n=36) from our dataset.

This filtering resulted in our fire dataset containing 641 fires across the five-year period (Appendix B). While the year with the most fires was 2023 (n=309), the year with the largest fires (those burning ≥ 100,000 acres) was 2020 (n=20 out of 33 fires). Then, we divided the size in acres by the duration in days for each fire to determine a daily estimate of acres burned across an individual wildfire’s lifespan. Finally, we summed across all active fires on a given day in order to calculate the estimate of daily acres burned by all fires in our study region (i.e., “wildfire acres”) (Kittelberger et al. 2022).

#### 2.3.3 Environmental Data

For our other environmental variables, we gathered data on precipitation (mm), minimum temperature (in degrees C; hereafter referred to as T_min), and drought index. We used minimum temperature in our analyses rather than another metric for temperature since it is likely a more accurate representation of physiological limits in birds when considering avian responses to climate change (Newton 2007, Tonelli et al. 2024). While our drought index data is reflective of a statewide average for Utah, the values for precipitation and T_min are specific to the Rio Mesa area. However, we would expect that temporal trends in these latter two variables are reflective of trends happening at larger spatial scales within western North America.

We downloaded daily precipitation and T_min data from the KNMI (2022) Climate Explorer covering 2011 through 2024 from the Glade Park meteorological station in neighboring Colorado (21 km away from the station). There was one day in our study (10/23/2020) for which there was no available climate data in this Glade Park dataset, so we averaged the precipitation and T_min data according to the values for the day before and after this date. For drought, we downloaded weekly categorical percent area drought index data for Utah from the US. Drought Monitor (Tinker et al. 2025) for the study period. We chose to use values for the “D2” drought category, which indicates “severe drought” (Tinker et al. 2025), in our analyses since a majority of weeks across the five years in our study period had non-zero values for this category within the drought index data. Additionally, utilizing the D2 data allowed us to test the impacts of more detrimental levels of drought on birds. For each day within a given week, our drought variable represented the D2 drought index for that week.

### 2.4 Statistical Analyses

For our analyses of banding data, we filtered out any records of birds that were not identified to the species level. We then created a series of linear mixed-effects model (LMEM) using the “lmer” function from the R package *lme4* (Bates et al. 2015) to test for the effects of wildfires on bird movements and health across years. For our environmental data, we first log-transformed our daily wildfire acre data. We then centered and scaled the rest of our environmental variables.

In our first model, we used abundance at our station based on daily captures as both a proxy for movement and our response variable. In addition to our environmental variable fixed effects, we also included year and Julian day as a fixed effect interaction term to account for variations over time. We also included the number of daily net hours as a random effect to account for the differences in sampling effort. For our second model, the body mass index was first calculated as the ratio of an individual bird’s mass at capture with the average mass for that species (Kittelberger et al. 2022), using average mass data for that species from the global avian trait database BIRDBASE (Şekercioğlu et al. 2025). We then used the body mass index as our response variable and again included our environmental variables and year-Julian day interaction term and also fat score as our fixed effects; for the latter, we included fat score in order to account for the effect of how much fat a bird has on its body mass (i.e. fatter birds should be heavier and have a higher mass index).

While our first model was a LMEM, we employed a phylogenetic generalized linear mixed-effects model (PGLMM) for our body mass index and body condition models to account for potential phylogenetic non-independence on avian body mass in our data. First, we downloaded 500 phylogenetic trees from http://birdtree.org (Hackett All Species; Jetz et al. 2012) for the bird species in our mass index and body condition datasets and created consensus trees using the “ls.consensus” function from the R package *phytools* (Revell 2012). Next, we used the “pglmm” function from the R package *phyr* (Li et al. 2020) with a Gaussian framework. Additionally, we included species as a random slope for year to allow the effects of time to vary by species.

Furthermore, in order to best account for the impact of lag effects of wildfires and other environmental variables on avian mass on the day a bird is captured (i.e., more accurately reflecting the temporal window in which environmental variables may be most influencing the fat deposition and body mass of a bird), we had our variables reflective of environmental conditions 7 days prior to each capture day. This one-week lag effect follows findings by Kittelberger et al. (2022) that looked at which models with various lag effect temporal windows best informed the bird data. We did not include a lag effect in the environmental variables for our bird abundance analyses since it has been shown that bird movements are more impacted by recent and proximate prior factors (e.g., Streby et al. 2015, La Sorte et al. 2018, Van Den Broeke and Gunkel 2021), and previous analyses showed that bird capture-wildfire models without lag effects performed better than those with lag effects (Kittelberger et al. 2022). For more information about accounting for lag effects with wildfires, see Kittelberger et al. (2022).

For both of our models, we tested for potential variance inflation factors (VIFs) among our fixed effects with the “vif” function from the R package *car* (Fox and Weisberg 2019). All VIFs were around or below 3 (Zuur et al. 2010). All statistical analyses and graphing were conducted in R (version 4.3.1, 2023-06-16; R Core Team, 2023).

### 2.5 Feather Preparation and Isotopic Analyses

For our stable hydrogen isotope (δ^2^H) analyses, we selected feathers from four species captured at Rio Mesa in which individuals showed signs of emaciation in the fall of 2020 (Kittelberger et al. 2022): Western Warbling Vireo (*Vireo swainsoni*), Gambel’s White-crowned Sparrow (*Zonotrichia leucocphrys gambelii*), Wilson’s Warbler (*Cardellina pusilla*), and Western Tanager (*Piranga ludoviciana*) (Figure 1). We selected feathers from individuals captured between 2019 and 2021 to compare birds before, after, and during the 2020 wildfire season. Overall, we had feathers sampled from 155 individual birds.

**FIGURE 1.**
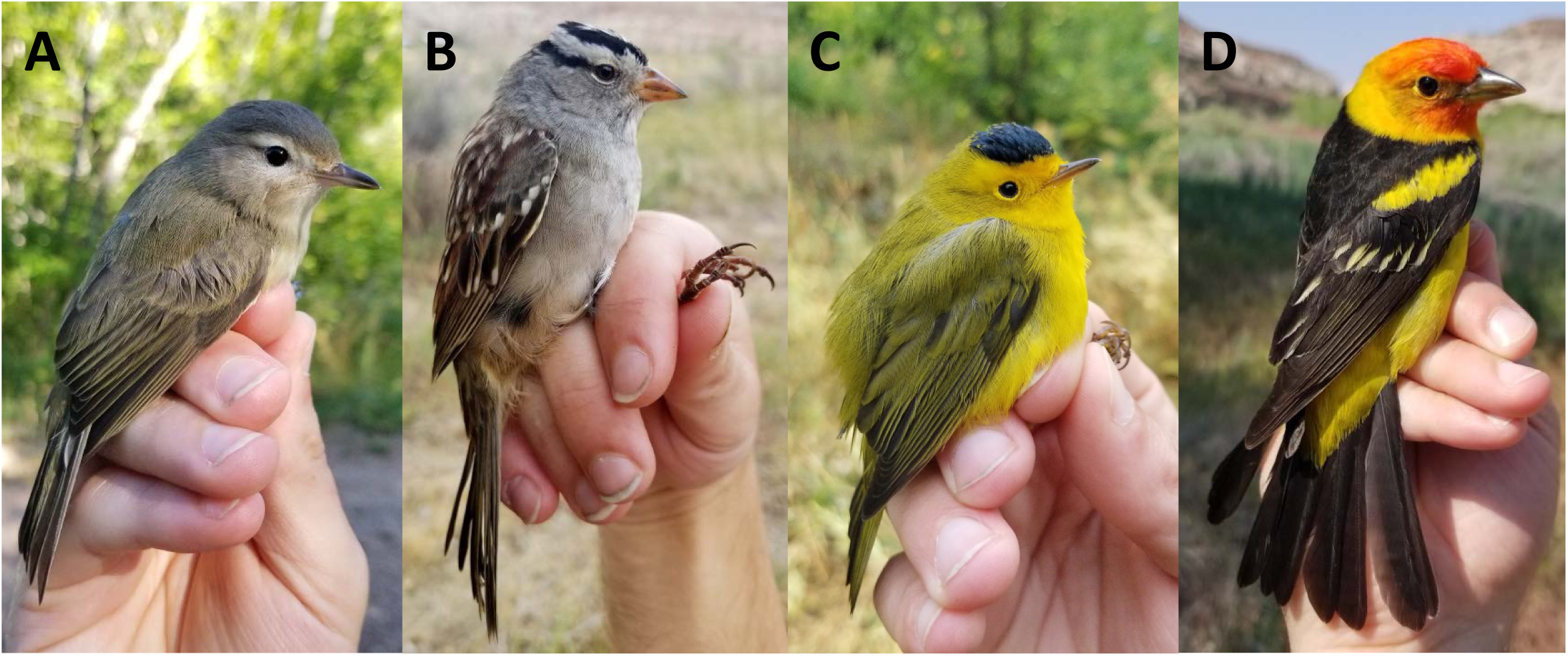
Focal bird species from which feathers were collected from Rio Mesa, Utah that were used for the hydrogen isotope analysis component of this study. These species showed heightened levels of emaciation and lethargy compared to other species banded during the fall of 2020 (Kittelberger et al. 2022). **A**) Western Warbling Vireo (*Vireo swainsoni*), banded on 8/25/22; **B**) Gambel’s White-crowned Sparrow (*Zonotrichia leucocphrys gambelii*), banded on 9/22/20; **C**) Wilson’s Warbler (*Cardellina pusilla*), male banded on 9/15/22; and **D**) Western Tanager (*Piranga ludoviciana*), male banded on 5/21/23. **A** & **C** taken at Red Butte Canyon Natural Research Area in Utah. All photographs taken by the lead author.

At the University of Utah SIRFER (Stable Isotope Ratio Facility for Environmental Research) laboratory, we first prepared feathers by cutting the distal third to half of a tail feather to use for isotopic analyses. We then created a 2:1 ratio of chloroform and methanol, respectively, by volume to effectively clean potential lipids from each sample (Norris et al. 2006, Magozzi et al. 2020). For a more detailed account of the feather processing protocol, see the Supplemental Material (Appendix C).

After ensuring all the methanol had evaporated from our samples, we then processed them for isotopic analyses. Using a razor blade, we cut each sample into small pieces and weighed them inside pre-baked silver foil capsules to a mass around 150 μg (ranging from 100-175 μg). We chose not to ground the feathers because of the potential for losing much of a sample due to the static nature of feathers. To account for the potential influence of the local atmosphere on hydrogen and to ensure the accuracy of isotopic results, a replicate was weighed for each sample. Reference standards with known δ^2^H values of non-exchangeable H were then incorporated into sample trays (Magozzi et al. 2020): Oryx antelope horn (ORX): δ^2^H = −34.0‰, δ^18^O = 25.09‰; Dall sheep horn (DS): δ^2^H = −172.7‰, δ^18^O = 6.02‰; and powdered keratin, POW: δ^2^H = −100.9‰, δ^18^O = 12.34‰. Capsules were then vacuum dehydrated for at least 7 days prior to isotope analysis (Magozzi et al. 2020). Samples were next pyrolized at 1400°C in an oxygen-free environment, and the resulting H_2_ and CO gases were then chromatographically separated and analyzed with a Delta Plus XL isotope ratio monitoring mass spectrometer (IRMS; Thermo Fisher Scientific) (Magozzi et al. 2020). Analytical uncertainty for the analyses, determined from the quality control POW reference, was respectively 1.9‰ and 0.1‰ (1σ) for the δ^2^H and δ^18^O values. For more background on these methods, see Magozzi et al. (2020).

Replicate analyses with tail feathers were run for samples (n=22) that either had δ^2^H values that may have been outliers (quite light or heavy) for their species, due to the possibility that these feathers may have been molted during migration and therefore not have originated on the summering grounds (Pyle et al. 2018), or if there was an issue with analyzing a prepared capsule initially (n=4). If the re-run values were heavier than the original ones, then we chose to stick with the original values. If the re-run data were lighter (and usually notably better and more in line with what we would have expected), we chose to use these newer values instead; we assumed that lighter values, regardless of the difference with the original-run result, were a more accurate indication of the geographic origin of the bird since these lighter values strongly correlate with increased estimated distance from origin to banding site. To ensure a standard approach across our data, we only used isotope data from rectrices, since body feathers and rectrices may not be molted at the same time or at the same location.

### 2.6 Analyzing Stable Isotope Data

In order to assign a geographic origin for our study individuals of unknown origin that were captured during migration, we utilized the R package *assignR* (Ma et al. 2020) to compare our isotopic values from samples of unknown origin with those with known origin. While *assignR* already has a library of known origin deuterium values for a variety of bird species, including Wilson’s Warbler, we searched for additional, available known origin reference data from other studies. We first focused our search on our four study species (and, in the case of White-crowned Sparrow, the Gambel’s subspecies specifically) (Paxton et al. 2007, Hobson et al. 2015, Ruegg et al. 2017, Lisovski et al. 2019) and then on congeners if there was no published isotopic data for our species of interest. We collected isotope data as well as corresponding geographic data and calibration methodology (Magozzi et al. 2021) in these publications for birds that were only described as being of known origin (e.g., from breeding sites). We then used the “refTrans” function to ensure that all reference data was on the same calibration scale as that used at SIRFER (“UT_H_1”) and used the “calraster” function to develop a feather δ^2^H map from the known origin data for each species (Ma et al. 2020).

We then generated probability density maps of unknown origin isotope data from Utah using the function “pdRaster”. For each species, we restricted the raster plots to a species’ breeding range. To do this, we downloaded eBird range GeoPackage files for our four target species (Fink et al. 2025) and then filtered these shapefiles to only focus on the breeding range for each bird. For Wilson’s Warbler, we edited the breeding range further to just focus on the area occupied by the “Western Boreal” breeding population as defined by the Bird Genoscape Project (Ruegg et al. 2014, 2020), as all but one of our warblers that had feathers analyzed for isotope analyses were also separately analyzed genetically and had been found to all belong to the Western Boreal group. We also restricted the eBird Warbling Vireo range to that representing the breeding area of the now recognized Western Warbling Vireo. Likewise, for Gambel’s White-crowned Sparrow, we adjusted the White-crowned Sparrow breeding range to only reflect that of the Gambel’s subspecies (Thorup et al. 2007, Welke et al. 2021).

We then visually inspected the generated maps for each bird to look for plots that seemed atypical (e.g., a plot showing a high probability origin south of Rio Mesa due to a higher than expected δ^2^H value; Figure S1) and to remove these from our study (n=2 for both the vireo and tanager). Like in other western songbirds, some Western Warbling Vireos and Western Tanagers are molt-migrants and replace some of their feathers at stopover sites (Pyle et al. 2009, 2018, Tonra and Reudink 2018, Kittelberger et al. 2025). Therefore, atypical values could reflect a feather that was grown the previous year during migration and not on the breeding grounds. This in particular has been an issue in prior studies that included samples from vireo species in the West, resulting in those deuterium results being excluded from the final analyses (Nordell et al. 2016). We also removed one Wilson’s Warbler sample from our analysis that was from a bird captured on 8/16/21, since this date was two weeks prior to our main study period. We did, however, include four samples of birds (warbler and vireo) from 2021 and one from 2019 that were collected up to four days prior to 8/31, due to both the availability of samples of the target species (i.e., to increase the sample size) and these occurring close to the start of the study period. Our final isotope dataset consisted of samples from 150 individuals across the four species.

For additional information about collecting isotopic reference material and importing it into *assignR* as well as creating these probability density raster plots, please see the Supplemental Material. All statistical analyses and graphing were conducted in R (version 4.3.1, 2023-06-16; R Core Team 2023).

### 2.7 Interpreting Isotope Data

In order to assess the impact of each fixed effect on the response variable, we first ran an analysis of variance (AOV) between the δ^2^H values and fixed effects for all species and an interaction term between year and Julian day, treating individual years as covariates. For heuristic purposes, we then assessed species separately, running an AOV for each between the δ^2^H values and an interaction term between year and Julian day.

Finally, since the δ^2^H isotope values themselves are not curtailed to a species’ breeding range, we used the function “wDist” from *assignR* (Ma et al. 2020) to calculate an estimated distance (utilizing a bird’s probability density plot) that each bird could have traveled from a probable origin to the Rio Mesa banding station. We chose to select the “w90Dist”, or 90% estimated cumulative distance, to best represent how far an individual bird could have traveled and converted our results into kilometers. We then ran an additional AOV for each species between the weighted 90% distances and the interaction terms between year and Julian day.

In order to help visualize the relationship between these sets of covariates, we created separate Non-metric Multidimensional Scaling (NMDS) plots using the R package *vegan* (Oksanen et al. 2025) for both δ^2^H and w90Dist, which also both contained Julian day and year; if we had species with less than four points for a given year, we created convex hull polygons rather than NMDS ellipses.

## 3. RESULTS

### 3.1 Environmental Trends

Among our four environmental variables (T_min, precipitation, drought, and wildfire acres) (Figure S2A, Table S1), we found significant positive trends across the five-year study period for only wildfire acres (coefficient: +0.050 ± 0.024 SE, t = 2.085, p = 0.038). This trend indicates that western North America has seen an increase in the amount of acres burned by wildfires. We found no significant trend in minimum temperature (+0.273 ± 0.229 SE, t = 1.191, p = 0.234) or precipitation (+0.162 ± 0.150 SE, t = 1.082, p = 0.280) around Rio Mesa, nor with severe drought in Utah (+1.123 ± 0.628 SE, t = 1.786, p = 0.075) during our study period.

### 3.2 Environmental and Banding Data Analyses

In our abundance model (Table S2), we found a significant positive effect of wildfire acres on daily bird captures across the study period (+4.712 birds/day ± 0.794 SE, t = 5.934, *p* < 0.001; Figure 2), indicating that more birds moved through the Rio Mesa site when there were a larger number of estimated daily acres impacted by wildfires in western North America. We found no significant effect of T_min, precipitation, drought, or the interaction between day and year on the daily number of captures between 2019 and 2023 (Figure 2).

**FIGURE 2.**
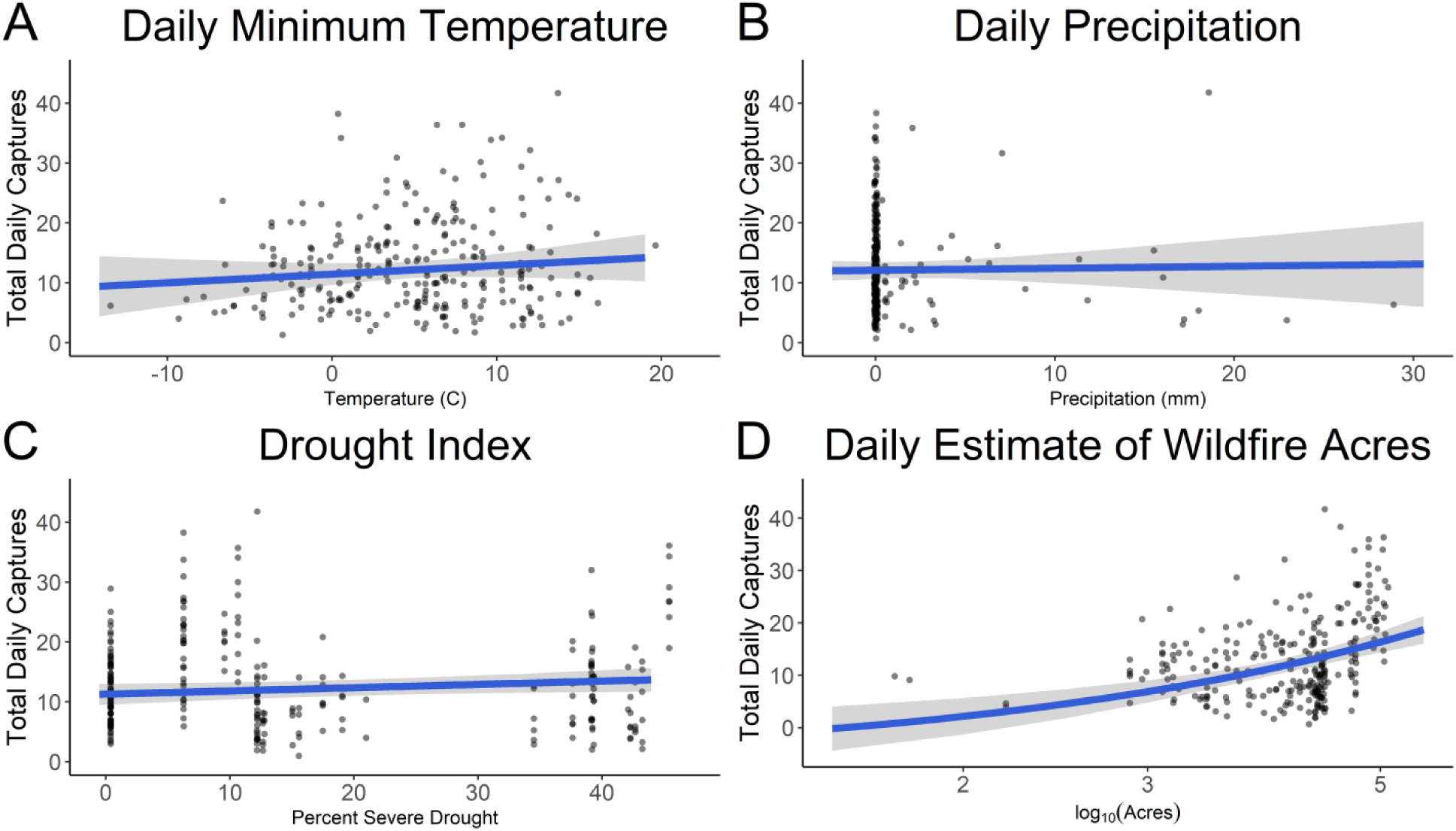
Trends in the daily number of birds captured at the Rio Mesa banding station in Utah (2019-2023) according to environmental variables: (**A**) minimum temperature, (**B**) precipitation, (**C**) severe drought, and (**D**) estimated wildfire acres. Results come from a linear mixed-effects model (Table S2), and the 95% confidence intervals are shown for each plot.

For our mass index PGLMM (Table S3), we found a significant negative effect of lagged (i.e. one week prior to capture) wildfire acres on bird weight across the study period (−0.016 ratio ± 0.002 SE, z = - 6.692, *p* < 0.001; Figure 3), indicating that individual birds captured at the Rio Mesa site were less heavy (i.e., more under the average weight for that species) when there was a larger number of estimated daily wildfire acres in western North America a week prior to capture. We found no significant effect of lagged T_min, precipitation, drought, or the interaction between day and year on the daily number of captures between 2019 and 2023 (Figure 3). As expected, there was a significant positive effect of fat score on the mass index (+0.023 ratio ± 0.001 SE, z = 21.589, *p* < 0.001), meaning that birds with higher amounts of body fat were heavier.

**FIGURE 3.**
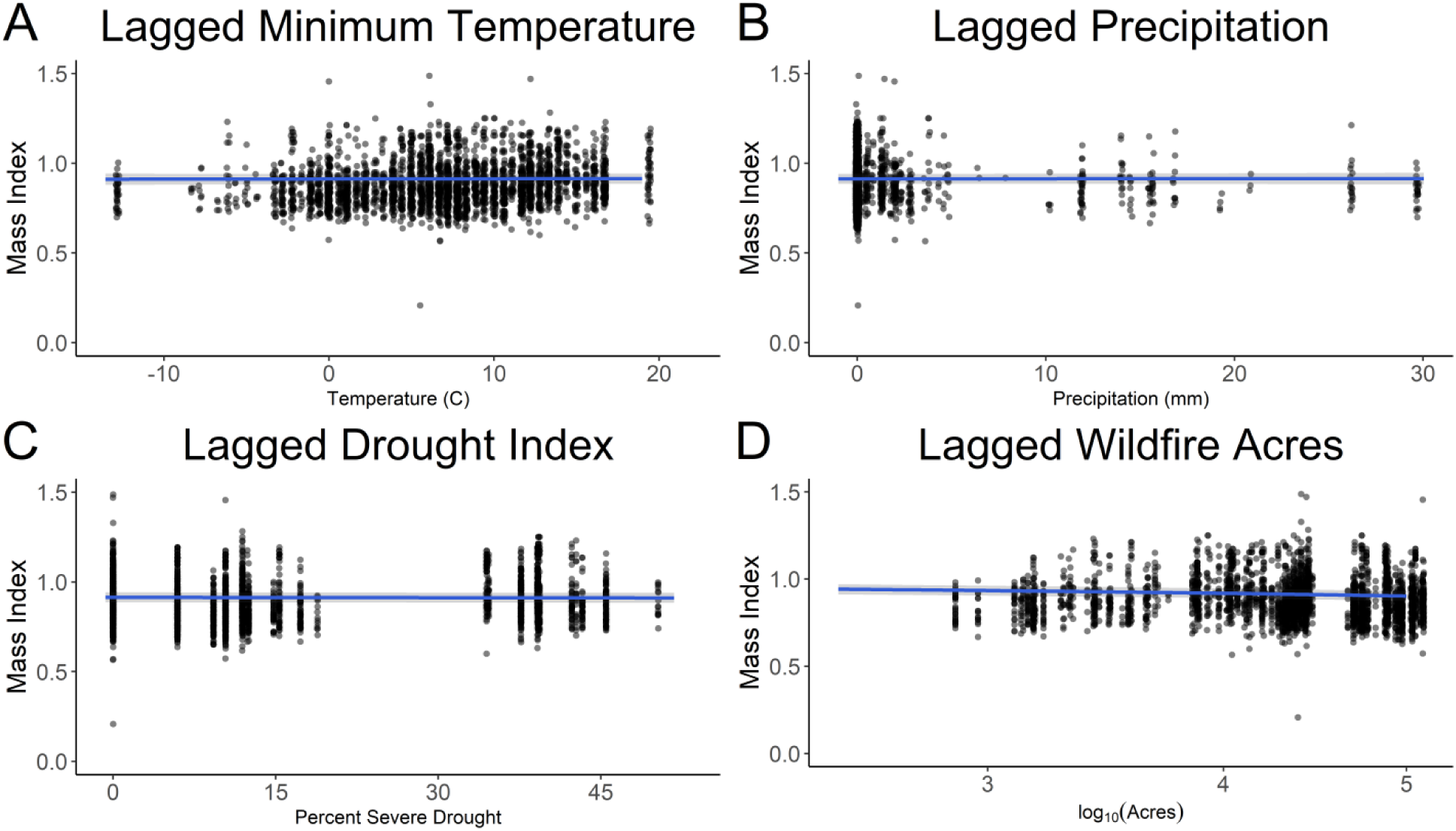
Trends in the mass index of individual birds captured at the Rio Mesa banding station in Utah (2019-2023) according to environmental variables that were lagged at one week prior to capture (see Methods): (**A**) daily minimum temperature, (**B**) daily precipitation, (**C**) weekly severe drought, and (**D**) daily estimated wildfire acres. Results come from a linear mixed-effects model (Table S7)^1^, and the 95% confidence intervals are shown for each plot. ^1^ While the statistical results noted in this paper for Mass Index come from a PGLMM, in order to be able to visualize these trends, we used a LMM instead, which did not produce qualitatively different results.

For our body condition model (Table S4), we found a significant positive effect of lagged drought (+0.015 score ± 0.006 SE, z = 2.453, *p* = 0.014; Figure S3), indicating that individual birds had a better body condition when there was more severe drought. We also found a significant negative effect of lagged minimum temperature on body condition (−0.016 score ± 0.008 SE, z = −1.975, *p* = 0.048; Figure S3), indicating that individual birds had worse body condition with warmer temperatures. In contrast to the trends for captures and mass index (Table S2,S3), there was no effect of year and/or Julian day on body condition (Table S4).

### 3.3 Stable Isotope Analyses

Our AOV of our full dataset of isotope values showed that both Julian day (*p* < 0.001) and species (*p* < 0.001) had significant effects on the δ^2^H values (Table S5; Figure S4A-B). There was no significant effect of year or the interaction between year and day on δ^2^H (Figure S4C). This indicates that while there was no significant shift over time in δ^2^H (outside of the expected daily trend within a season when birds from different locales tend to migrate on different days as fall migration progresses (Paxton et al. 2007, Conklin et al. 2010, Neufeld et al. 2021)) and thereby geographic origin of birds at the community-level, there was a notable effect of species (particularly Western Warbling Vireo and Western Tanager) on the isotope values (Figure S4A-B).

At the species level, we found that in AOV’s for all four species (Table S5), there was no significant effect of year, Julian day, or the interaction between these two terms on the δ^2^H values. However, for Western Warbling Vireo, the effect of the interaction between year and day was near significant (*p* = 0.066). Looking at the NMDS plots for all four species, we can see that in both Western Warbling Vireo and Western Tanager, δ^2^H values of birds from 2021 clustered away from the other two years (Figure 4A,D), while there was much overlap in the samples for birds across all three years for both Wilson’s Warbler and Gambel’s White-crowned Sparrow (Figure 4B-C).

**FIGURE 4.**
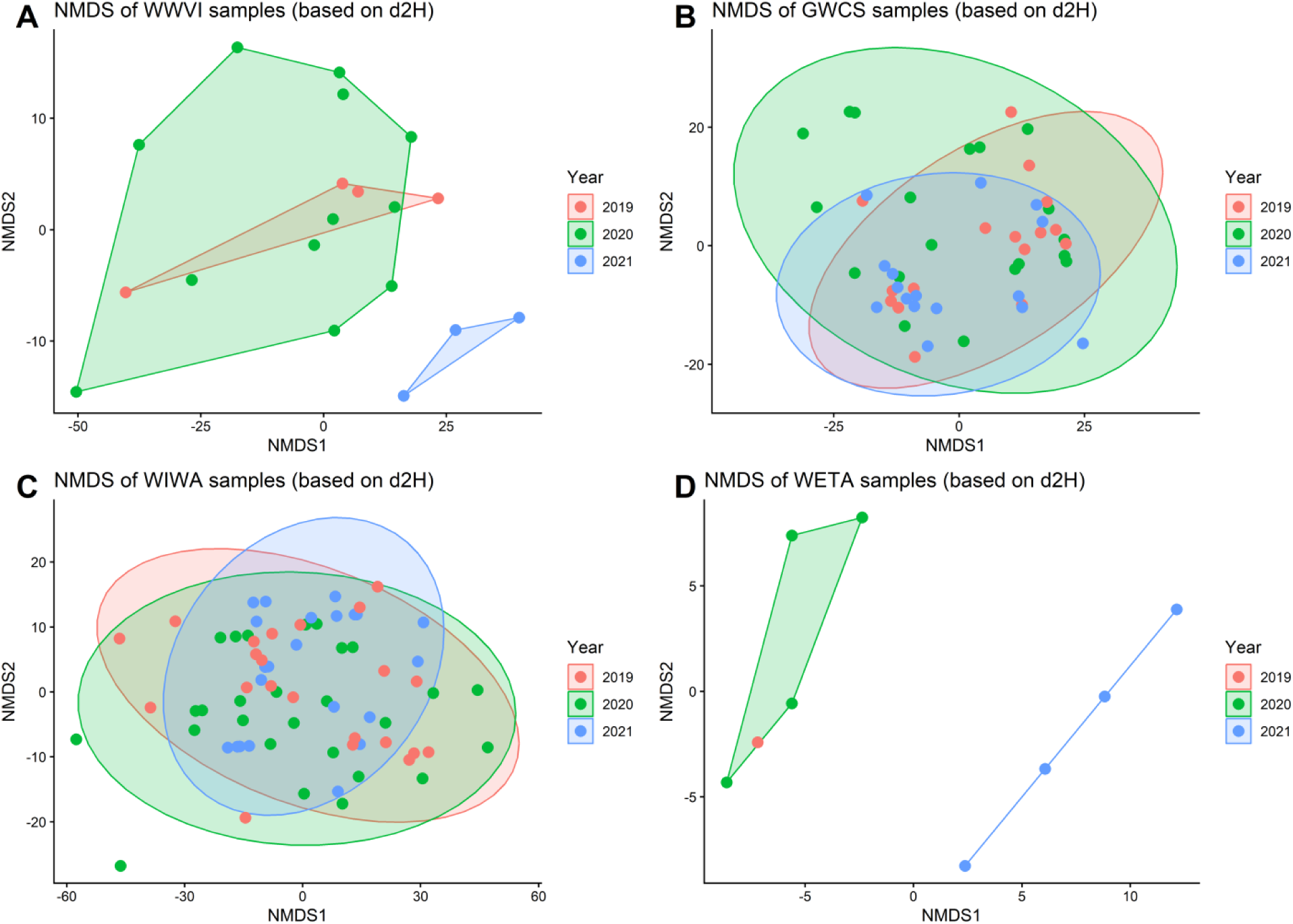
Non-metric Multidimensional Scaling (NMDS) plots between δ^2^H isotope values, Julian day, and year for four species of birds: **A**) WWVI = Western Warbling Vireo, **B**) GWCS = Gambel’s White-crowned Sparrow, **C**) WIWA = Wilson’s Warbler, and **D**) WETA = Western Tanager. For species with fewer than four points for a given year, convex hull polygons rather than NMDS ellipses are depicted. See Table S5 for results from analyses of variance between these variables for each species.

For our second set of AOV’s for all four species (Table S6), we found that there was no significant effect of year, Julian day, or the interaction between these two terms on the weighted 90% distances. However, again for Western Warbling Vireo, the effect of the interaction between year and day on distance was near significant (*p* = 0.060). Looking at the NMDS plots for all four species, we can see that in both Western Warbling Vireo and Western Tanager, estimated distances traveled for birds from 2021 clustered away from the other two years (Figure S5A,D), with 2021 birds arriving earlier and in general from closer origin sites (Figure 5A,D). For both Wilson’s Warbler and Gambel’s White-crowned Sparrow (Figure S5B-C), there was much overlap in the samples for birds across all three years, meaning that birds in general were traveling similar distances and therefore were likely from similar origin sites in all three fall seasons (Figure 5B-C).

**FIGURE 5.**
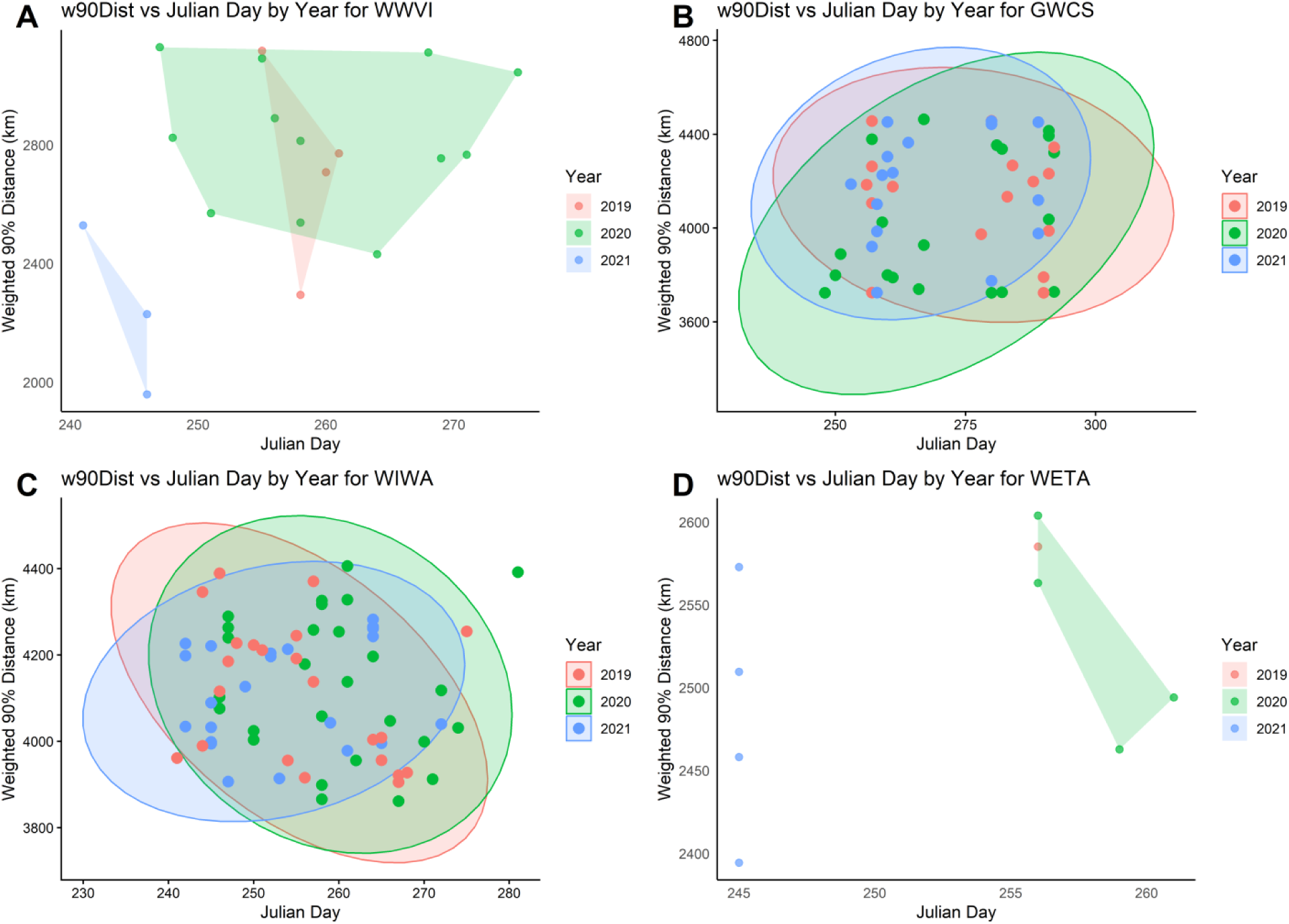
The distribution of weighted 90% distance estimates (“w90Dist”) over time by Julian day and year (2019-2021) for four species of birds: **A**) WWVI = Western Warbling Vireo, **B**) GWCS = Gambel’s White-crowned Sparrow, **C**) WIWA = Wilson’s Warbler, and **D**) WETA = Western Tanager. For species with less than four points for a given year, convex hull polygons rather than ellipses are depicted.

## 4. DISCUSSION

In this study, we used a multifaceted approach involving multiple years of bird banding and hydrogen isotope data from a bird banding station in southeastern Utah in order to assess how wildfires may be impacting bird communities during fall migration in the western United States. Previous analyses of fall 2020 banding data from this station found that higher daily bird captures were correlated with more acres burned by proximately-occurring wildfires (Kittelberger et al. 2022). Here, we confirmed this relationship across our five-year study period (Figure 2). We found that not only was there a significant positive effect of day-of wildfire acres on daily bird captures, but also that there was no significant effect of temperature, precipitation, or drought on these daily captures (Figure 2, Table S2). This means that more birds were moving through the region of our banding station when wildfires were concurrently more active. Likewise, this finding emphasizes that, at least in recent years, more active and impactful wildfires are likely to have a more important influence on bird movements across the landscape (Overton et al. 2022, Stanek et al. 2025) and therefore on abundances at stopover sites than do other environmental variables (at least, the major ones that we included in our study) during fall migration in western North America (Figure 2). Our results are likely strongly affected by the fires in 2020, and we could expect that in a year with a non-severe wildfire season, other environmental variables would play a more important effect on birds.

We also found a significant negative effect of wildfire acres one week prior to banding, on our body mass index. Again, there was no significant effect of temperature, precipitation, or drought on body mass index (Figure 3, Table S3). This significant effect of lagged wildfire acres on mass mirrors what we found previously in 2020, with reductions in body mass of birds correlated with more wildfire acres one week prior to capture (Kittelberger et al. 2022). We reaffirm this relationship after simultaneously testing the influences of other environmental variables and accounting for the expected influence that changes in fat loads and date have on mass throughout the season on migratory birds (Table S3). In fact, the decline in body mass that we found in birds across the five year study is reflective of a larger trend found across thirteen years of banding data from this station (Kittelberger et al. (2025) *in review*), though the specific reason(s) for this are not yet clear. Here, we show that when there are more acres affected by wildfires during or before a bird’s journey, birds tend to have lower body masses, showing how wildfires can adversely affect birds during an already risky and energetically demanding period of the year. There is a number of potential reasons why wildfires could cause birds to lose body mass, such as forcing them into atypical or more competitive habitats, causing them to fly further and outside of their expected path, not being able to rely on prior stopover sites due to the sites being burnt, or skipping stopovers altogether to continue flying and therefore not being able to refuel due to fire or smoke (Frost 2018, Overton et al. 2022). While we did not account for smoke particulate matter in this study, prior research has shown that when air quality is worse due to wildfire smoke, the body mass of birds declined, perhaps because birds must expend more energy in such adverse conditions (Nihei et al. 2024).

On the contrary, we found that there was no significant effect of lagged wildfire acres on body condition, and instead birds had better condition with more severe drought and worse condition with warmer temperatures (Table S5). At first, these results may seem both contradictory with the effect we found of wildfires for body mass and also counterintuitive with expected trends for these environmental variables. However, our body condition model only uses data beginning in 2021 and therefore does not include data from the fall 2020 wildfire season, since this metric was developed to better assess an individual bird’s health in response to our observations of emaciated birds in the field during 2020 (Kittelberger et al. 2022). Therefore, body condition data only began to be fully recorded in 2021, and there also has not been as severe of a wildfire season as 2020 since then. In fact, 2023 was the weakest season in 25 years in terms of acres burned, even if there have been more individual fires in some of the years that followed (NIFC 2025). We are thus not surprised to see a lack of effect of wildfire acres on birds during 2021-2023, and likewise have not seen the heightened levels of emaciation and poor body condition on birds that we saw in 2020, positive indicators for the health of migratory birds in recent years. With regards to drought (Figure S3), while we would expect to see birds with worse body condition under drier conditions, a prior study showed that birds in Utah state tend to concentrate in riparian habitats due to resource availability when conditions are drier (Neate-Clegg et al. 2021), and droughts can drive large shifts in the abundance of bird communities (Albright et al. 2010). Therefore, drought could actually be forcing birds into areas with better vegetation quality and more food, especially in wetter monsoon years (Stanek et al. 2025), thereby leading to better rather than worse body condition. As for the negative effect of increasing minimum temperature on body condition (Figure S3), warmer conditions could cause heightened levels of thermal stress (e.g., dehydration) on birds (Wolf 2000, Albright et al. 2017), especially when moving through an already warm and dry region that is a climate change hotspot (Allocca et al. 2025), and thus result in worse body condition (particularly in young birds) (Oswald et al. 2021). We recommend that banding stations continue or begin to record body condition data in order to better assess the health of birds.

While we did not examine the actual mechanisms by which birds respond to wildfire activity and instead focused on the responses themselves with our banding data, we used heavy hydrogen isotopes to try to better understand how wildfires may have influenced from where and when birds were migrating. We first found that day of capture (serving as a proxy for day of departure, i.e. later capture days could suggest later departures dates) had a significant effect on the isotope values of our bird community (Table S5). This is not surprising, as birds migrating at different periods within a season are likely originating from different regions and/or latitudes (Paxton et al. 2007, Conklin et al. 2010, Neufeld et al. 2021). However, we did not find any significant indication in our models that estimated migratory origin or distance traveled to our banding station were shifting over time between 2019 and 2021 (Table S5, S6). This would suggest that in terms of where birds were migrating from, nothing major changed for these species in response to wildfires. However, our day of capture variable may not actually be a good proxy for departure date, especially as we are not able to determine how long individual birds were migrating. For instance, while the origins and estimate distances of different birds may ultimately be similar, we do not know the paths birds took during migration, where they stopped over (or the number of stopovers taken), or how the duration of migration varied between years, all of which can be influenced by fire activity (Overton et al. 2022). Our estimated distance traveled variable is also a simplification of the more nuanced movements of birds that are not necessarily in a straight line (see Overton et al. (2022) for a great example of migratory paths that had the same endpoints but differed widely between individuals and across years).

Additionally, all of the feathers in this study come from different individuals (i.e., no sampled birds were recaptured in a different year during the study), meaning we were not able to examine the feather isotopes of the same bird over time to look for any spatio-temporal shifts at the individual level. Finally, our analyses are limited by our sample sizes, reducing the measured magnitude potential trends. In fact, a power analysis on a linear model with 2 predictor variables (day and year), assuming a strong effect size (f^2^ = 0.35), would require 27-28 individuals in order to see a significant difference, which is more than the number of vireo and tanager samples in our dataset. We did have to remove some individuals from our dataset that had feathers likely molted off the summering grounds, though this was a relatively small number. These feathers mostly belonged to known molt-migrant species, also meaning that there is chance that the δ^2^H values from the remaining included feathers for these species may reflect molting grounds rather than more northern summering grounds (which could influence estimated distances and our models), even if the δ^2^H values suggest a plausible area within the species’ expected breeding range. While we strove to use a feather from every banded individual of the target species across the study period, the reality is that in 2019 and 2021, there were just far fewer captures of certain species.

We can actually see the effect of a smaller sample size on our results when looking at our NMDS plots. For two species, Western Warbling Vireo and Western Tanager, there were differences in both the δ^2^H values and estimated distances between years, with 2021 birds visibly clustering separately (in contrast with birds for the other two sampled species) (Figure 4,S5). Specifically, we see that vireos and tanagers in 2021 migrated earlier in the season and from seemingly overall closer locations compared with those from the previous two years (Figure 5). It is not exactly clear what changed for these two species between 2020 (and even 2019) and 2021, but there may have been a loss of individuals migrating through southeastern Utah originating from some areas and a gain of individuals from other sites (especially for the vireo). Some of the populations from prior breeding sites may have perished during or not fully recovered from the avian mortality events of 2020 or perhaps some populations initiated their migration earlier in 2021 in response to residual carryover effects from the prior year. In fact, about half as many birds migrated through the station in both the spring and fall seasons of 2021 compared with the fall 2020 season (authors’ unpublished data).

We also found in our models that that there was a near-significant interaction between year and day on both δ^2^H and distance for Western Warbling Vireo (Table S5,S6). This suggests that there was a shift in departure days or at least in migration timing across the three years for this species, a trend that may have become significant with a larger sample of individual vireos (either from other years or other sites). We can further see for the vireo how there was a major influx of individuals of this species in 2020, across a larger migratory window and from primarily further sites, compared to the pattern before and after this year, when vireos had a narrower window during which they were migrating through this particular stopover site (Figure 5A). This reinforces our findings from our abundance banding model (Table S2) and highlights how movements and abundances of this species were affected during a severe wildfire season. Interestingly, we do not see much of a difference in the likely origins of Wilson’s Warblers and Gambel’s White-crowned Sparrows across the three years (Figure 5,S4B). This could suggest that either the populations of these two species potentially recovered better (or were affected less) from the prior wildfire season compared to the vireos and tanagers that migrate through at least the Interior West, or that the movements themselves that the warblers and sparrows took in 2020 were not as influenced by fires (even if the condition of these species were (Kittelberger et al. 2022)).

More broadly, our isotope work increases our knowledge of the likely geographic origins of migratory birds in this flyway (Figure S6-10,A-C) and adds to the literature on feather-based hydrogen isotope analyses. For example, it is of note that all of the Western Tanagers in 2021 were migrating through and captured on the same day at Rio Mesa even though they seemingly originated from different locations, which may be a result of the species perhaps having a narrower migratory window in the region in the fall (and captures in 2020 spanned just several days) compared to other Neotropical migrants like Wilson’s Warbler that migrate over a more extended period of time (Figure 5). Additionally, our work effectively compiles in one place the previously published isotope data for Wilson’s Warbler and Gambel’s White-crowned Sparrows (Appendix D), which will be useful for other researchers studying feather isotopes for these or similar species. Notably, this is the first study to our knowledge that publishes δ^2^H data for Western Warbling Vireo and Western Tanager (Figure S6A-B,S8). While our estimated origins for these species are tentative due to these species being molt-migrants, we know that at least some of the Western Tanagers that migrate through Rio Mesa travel as far north as southwestern Canada for the summer, based on a Motus tagged individual from Rio Mesa (ID# 54664) that migrated northwards and was detected at two sites in southern British Columbia; this aligns with the broad likely origin region for most of our sampled tanagers in this study (Figure S1). Because there was no previously published known-origin data for the breeding individuals of these two species, and we had to instead rely on data from species that shared similar foraging strategies and breeding grounds, future isotope data from these species’ summering grounds will help refine the probability density plots for unknown origin migratory individuals. Moreover, known-origin isotope data will also importantly reveal how accurate the δ^2^H values of feathers collected during the summer are with where the feather was collected, and thereby help inform future isotope work for these species and other similar molt-migrants during fall migration.

## 5. CONCLUSION

In this study, we built on our prior work from the Rio Mesa banding station in Utah on the effects of wildfires on birds during fall migration (Kittelberger et al. 2022), by applying the previous framework of daily estimated wildfire acres to a five-year period rather than just 2020 and examining the effect of other important environmental variables concurrently against wildfire acres. We confirmed that when there are more wildfires active in western North America, there are more captures of birds in the Interior West, likely a result of birds shifting their movements to avoid areas of fire and smoke (Overton et al. 2022), and that birds have worse body conditions. Our study also indicates that during periods when wildfires are especially active and severe, wildfire activity has the most significant influence on bird movements and health (Figure 2,3), more so than other important environmental variables. The results from both of our body mass and body condition models also have important implications for understanding threats to migratory birds and their health in the future, especially against longer-term background changes in avian physiology (Kittelberger et al. 2025 *in review*), as wildfires are expected to continue growing in size and intensity in western North America in the future (Buechi et al. 2021). While we are not able to account for the effect of more impactful wildfires on body condition prior to 2021, the trends we did find for drought and temperature help us better understand how other environmental factors affect the health of birds in years when wildfires are not as severe. Moreover, the difference in trends we found between our mass index and body condition models support earlier work that showed droughts and wildfires can have differing effects on some bird communities (Stanek et al. 2025).

Additionally, the heavy hydrogen isotope component of this study allowed us to explore more than just the day of captured birds and to try to uncover more of the story by tracing individual bird migration back towards where birds were originating from. Specifically, we aimed to determine an estimated origin for birds during the summer on their breeding or post-breeding grounds and explore if birds were shifting their departures from certain areas over time. While we broadly did not necessarily see a shift in departure dates, we did see evidence that Western Warbling Vireo and Western Tanager varied across year in both their migration timing and likely summer origin, even with our sample limitations and lack of significance. For Western Warbling Vireo, we found a near-significant trend in migration dates changing across years and can see that there was a wider migratory window (in addition to more individuals captured) of birds in 2020 compared to before and after this severe wildfire season (Figure 5, Table S5,S6), emphasizing how bird movements were affected that year. We also note that both of these species show a notably different pattern in hydrogen isotope values in 2021 compared to the two previous years, which should be explored in future research. More broadly, our study contributes to the previously established hydrogen isotope bird feather literature for Wilson’s Warbler and White-crowned Sparrow (Figure S7,S9A-C) and notably introduces some of the first published δ^2^H data for the tanager and vireo (Figure S6A-B,S8). Combined with the code of our workflow in this study for analyzing hydrogen isotope data (Kittelberger 2026), our isotope data and code can aid other researchers in the field and provide a foundation for future work on examining the migratory origins of western North American birds, especially for our four focal species. Ultimately, our isotope work may reveal that the most important piece of the puzzle for understanding the effects of wildfires on migratory birds and their health (and one that hydrogen isotopes cannot necessarily inform about) is not a bird’s migratory origin but instead the period after migration has been initiated and before capture at a site like Rio Mesa, as longer continuous flights and fewer stopovers would provide birds with fewer opportunities to replenish fat reserves and thereby heighten emaciation risk.

## 6. ACKNOWLEDGEMENTS

We are grateful to Zachary J. Lundeen and Hau Q. Truong for their long-term friendship and assistance with seasonal operations on site at Rio Mesa. We thank the USGS Bird Banding Lab for their priceless contributions to ecological research over the years and for providing bands and banding permits to the authors, and the Utah DWR for providing a permit for our station. We are grateful to Suvankar Chakraborty and the SIRFER Lab for assisting with the isotope analysis component of this study. We thank Megan Miller, Lola Phillips, Nicholas Seefeldt, and Alexia Tzortzis for their involvement in either collecting wildfire data and/or assisting with preparing feathers for isotope analyses. We also are grateful to Kristen Ruegg, Amanda Carpenter, Christen Bossu, and the rest of the Bird Genoscape Project team for analyzing our Wilson’s Warbler feathers, providing us with those population-designations, and communicating about other ornithological topics, and to Marilyn Ramenofsky for helping provide the known-origin reference isotope data for Gambel’s White-crowned Sparrows. Finally, we thank all of the countless lead bird banders and banding assistants that have been a part of banding operations at Rio Mesa and helped collect some of the data used in this study. We are grateful for the generous support of H. Batubay Özkan and Barbara Watkins through the years.

## FUNDING

We thank the Redd Center at BYU (through their Summer Award for Upper Division and Graduate Students for 2021 and 2022), the University of Utah’s Bonderman Field Station at Rio Mesa, and the Medium Grant from the Sustainable Campus Initiative Fund (SCIF) at the University of Utah for helping fund and support operations and equipment used during banding seasons. We thank the Wilkes Center for Climate Science & Policy for helping fund the isotope analyses. We are grateful to the Graduate Research Fellowship and the Vincent Hillyer Fellowship at the University of Utah for their support of the lead author.

## 7. AUTHOR CONTRIBUTIONS

KDK conceived the original idea for the project; KDK oversaw and managed the field research component of this study, with support from ÇHŞ; KDK and LWS prepared the feathers for isotope analyses; KDK prepared the datasets; KDK statistically analyzed the data and created the figures, with assistance from GJB and CJT; KDK wrote the manuscript; all co-authors contributed to and gave final approval of the manuscript.

## 8. CONFLICT OF INTEREST

The authors declare that there are no commercial or financial relationships in this research that could be seen as conflicts of interest.

## 9. SUPPORTING INFORMATION

See the Supplemental Material for additional information. The various datasets, files, and code utilized and referenced in this study can be found on Zenodo (https://doi.org/10.5281/zenodo.18379587; Kittelberger 2026).

## Notes

https://doi.org/10.5281/zenodo.18379587

### Competing Interest Statement

The authors have declared no competing interest.

