## Supplemental Material for "Wildfires during fall migration affect bird movements, avian health, and migration timing in western North America: Insights from bird banding and feather isotopes"

### 1. Additional Methodology for Analyzing Stable Isotope Data

#### *1.1 Workflow for Incorporating Known-origin Isotope Values*

In order to assign a geographic origin for our study individuals of unknown origin that were captured during migration, we utilized the R package *assignR* (Ma et al. 2020) to compare our isotopic values from samples of unknown origin with those with known origin. While *assignR* already has a library of known origin deuterium values for a variety of bird species, including Wilson's Warbler, we searched the literature for additional, available known origin reference data from other studies. We first focused our search on our four study species (and, in the case of White-crowned Sparrow, the Gambel's subspecies specifically), resulting in us finding data for Wilson's Warbler and Gambel's White-crowned Sparrows (Paxton et al. 2007, Hobson et al. 2015, Ruegg et al. 2017, Lisovski et al. 2019). We coalesced this isotope data as well as corresponding geographic data (latitude and longitude, in decimal degrees), isotope uncertainty values (which were between 1.9 and 2.0), and calibration methodology following Magozzi et al. (2021). We only selected isotope data in these publications for birds that were described as being of known origin (e.g., from breeding sites) in order to ensure the sampled feathers were likely replaced on the breeding or post-breeding grounds, rather than at a migratory stopover site or on the non-breeding grounds. We then subsetted our data to that of the two species.

After correctly formatting our column names (see the included R script for this study), we used the "refTrans" function to ensure that all reference data was on the same calibration scale as that used at SIRFER ("UT_H_1"), resulting in hydrogen isotope values being accordingly update. We then reordered the columns in our dataframe to ensure that the isotope data (values and corresponding uncertainty) and geographic coordinates came first. Next, we converted our data into a vector of "SpatVector" class using the latitude and longitude of the isotope data, and downloaded a fine-scale water isoscapes using the "getIsoscapes" function, filtering to just the hydrogen isoscapes. We then used the "calraster" function to import the known origin data for each species into *assignR*, using the previously downloaded hydrogen isoscapes and masking the plots to North America. For our two species for which we could not find known-origin data (or that of congeners) in the literature, we used the library of data included in assignR; for Western Warbling Vireo, we used the function "subOrigData" to select data for White-eyed Vireo (*Vireo griseus*) and Red-eyed Vireo (*V. olivaceus*), while for Western Tanager we selected data from Swainson's Thrush (*Catharus ustulatus*) and Northern Cardinal (*Cardinalis cardinalis*) to best represent that species. We then followed the steps noted previously to prepare this known origin data (see the R script for more).

***1.2 Workflow for Analyzing Unknown-origin Isotope Values***

We then imported our unknown origin isotope data from our samples in Utah. For each species, we restricted the raster plots to a species' breeding range. To do this, we downloaded eBird range GeoPackage files for our four target species (Fink et al. 2025) and then filtered these shapefiles using ArcGIS Pro (ESRI 2021) to only focus on the breeding range for each bird. For Wilson's Warbler, we edited the breeding range further to just focus on the area occupied by the "Western Boreal" breeding population as defined by the Bird Genoscape Project (Ruegg et al. 2014, 2020), as all of our warblers that had feathers analyzed for isotope analyses were also separately analyzed genetically and had been found to all belong to the Western Boreal group. We also restricted the eBird Warbling Vireo range to that representing the breeding area of the now recognized Western Warbling-Vireo. Likewise, for Gambel's White-crowned Sparrow, we adjusted the White-crowned Sparrow breeding range to only reflect that of the Gambel's subspecies (Thorup et al. 2007, Welke et al. 2021). We used the function "st_read" to incorporate these GeoPackage files into R and converted them into a "SpatRaster" file using the function "vect". Due to the GeoPackage files not necessarily being of the same spatial extent as North America, we used the "project" and "rasterize" functions to align the breeding ranges onto our mean hydrogen isoscapes map for North American; we used the function “compareGeom” to check that the breeding range and isoscapes were correctly aligned. Finally, we generated probability density plots for our samples of each species using the function “pdRaster” and the previously generated hydrogen isoscapes for North America, restricting the plots to our species’ breeding range priors.

Finally, we used the function “wDist” from *assignR* (Ma et al. 2020) to calculate an estimated distance that each bird could have traveled from a probable origin to the Rio Mesa banding station. To do this, we provided the wDist function the probability density maps for each individual of a species as well as a spatial vector of the coordinates for the Rio Mesa station. While wDist provides a mean probable distance as well as multiple weighted probability estimates (all in meters), we chose to select the “w90Dist”, or 90% estimated cumulative distance, to best represent how far an individual bird could have traveled.

All statistical analyses and graphing were conducted in R (version 4.3.1, 2023-06-16; R Core Team 2023).

### 2. Study Design for Potential Future Sampling Protocol

A follow-up project to the current one could further use heavy hydrogen ($\delta^2H$) isotopes, from collected feathers to predict the likely known summer origin or estimated distance traveled of birds captured at a migratory stopover site(s), to further examine the influences of environmental variables (such as wildfires, droughts, etc.), temporal trends, or some other factor on bird migration and breeding. We recommend that such a study aim for around 28 or more samples for a species in a given year, in order to see clearly any potential significance at a “strong effect” level. If this number of samples is not possible for a given year from a single location, then we recommend that researchers pool samples from other study sites as well in order to hit the target power level. In this case, site can be treated as a random effect in statistical models or as a fixed effect, if there is a desire to test the difference in sites as well; the latter would likely be an important consideration if sampled sites are far apart and/or from different habitats/ecosystems. Note though that if site is treated as a fixed effect, then a larger number of individuals will be needed from separate sites in order to increase the power of analysis for difference among sites. For any species that is a known molt migrant, such as Western Warbling Vireo and Western Tanager in the current study, we advise researchers to pull more than one tail feather (and likely a different rectrix too; i.e., R4 on the left, R3 on the right) in case these feathers were grown at different locations and isotope values for one feather are outside of the expected range.

This kind of study protocol can be applied to samples from the four species included in the current study, or to samples from additional species. We recommend that researchers follow the workflow established before this section for working with known origin reference data and unknown origin field samples, and we emphasize that alternative references will be necessary for any species chosen that is either not already included in the *assignR* database or has not been previously sampled in the literature. Additional helpful information can be found in the methodology in the manuscript, as well as within our available R script.

**FIGURES AND TABLES**

**FIGURE S1.** Example probability density plots for four Western Tanagers across the species' breeding range, which were banded at the Rio Mesa banding station in Utah during fall migration in September, 2020. The first two individuals (#23,25) have plots that show estimated origins (darker green represents higher likelihood of origin) from the central to northern parts of the species' summer range, which would be expected for a bird migrating through Utah in the fall; this was the pattern for most of the tanagers sampled for this study. The bottom two individuals (#48,49) have plots that suggest the origin of analyzed feathers came from the southern tip of the breeding range (or even south of that). Since Western Tanager is a molt-migrant, this would indicate the feathers for these two birds were likely replaced previously at stopover sites rather than on the breeding grounds. These bottom two birds were subsequently removed from the final set of analyses in this study.

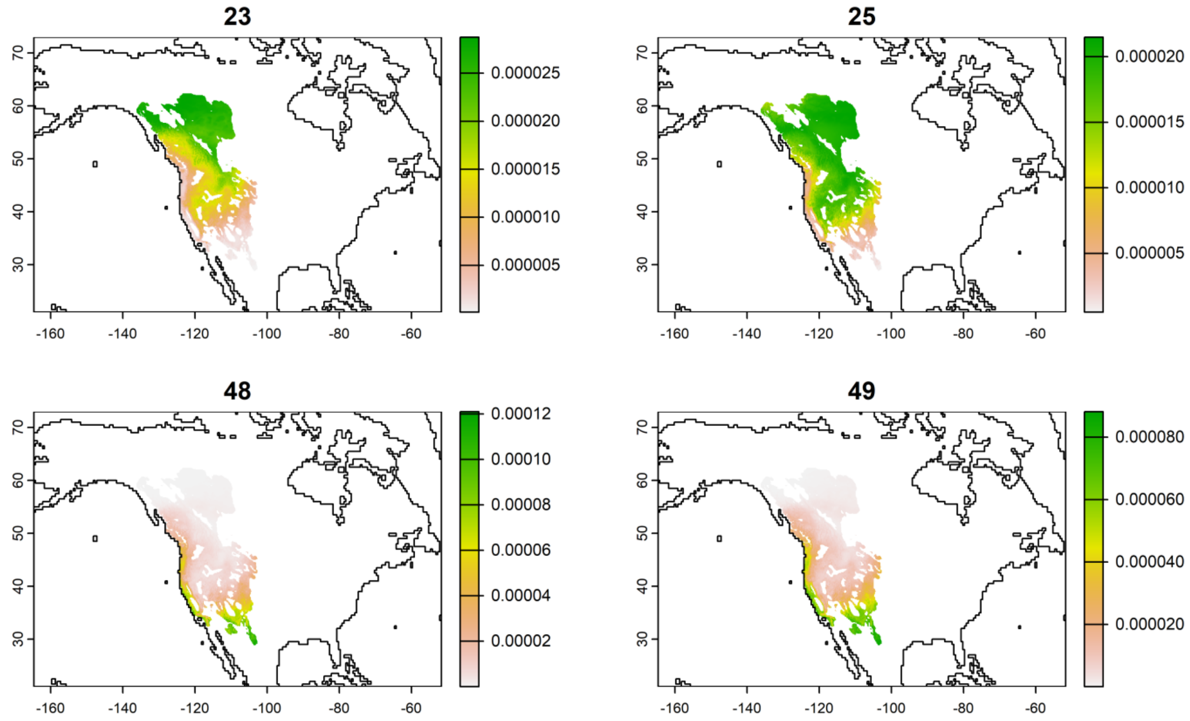

**FIGURE S2.** Temporal trends across our study period (2019-2023) in the fall for our environmental variables: (**A**) daily minimum temperature, (**B**) daily precipitation, (**C**) weekly severe Drought Index, and (**D**) daily estimate of wildfire acres. Trendlines are fitted from linear models with 95% confidence envelopes. See the Methods for information on the sources of and specifications for these various variables.

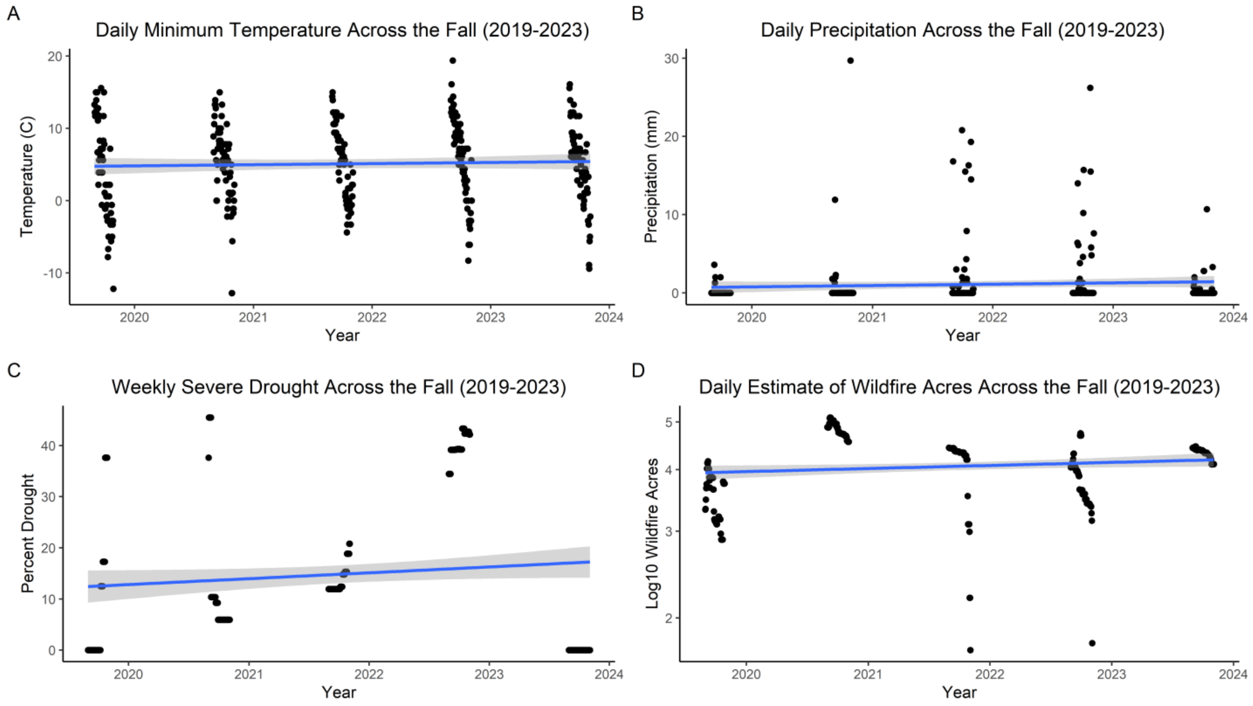

**FIGURE S3.** Trends in the body condition of individual birds captured at the Rio Mesa banding station in Utah (2021-2023) according to environmental variables that were lagged at one week prior to capture (see Methods): (**A**) daily minimum temperature, (**B**) daily precipitation, (**C**) weekly severe drought, and (**D**) daily estimated wildfire acres. Results come from a linear mixed-effects model (Table S8)[1], and the 95% confidence intervals are shown for each plot.

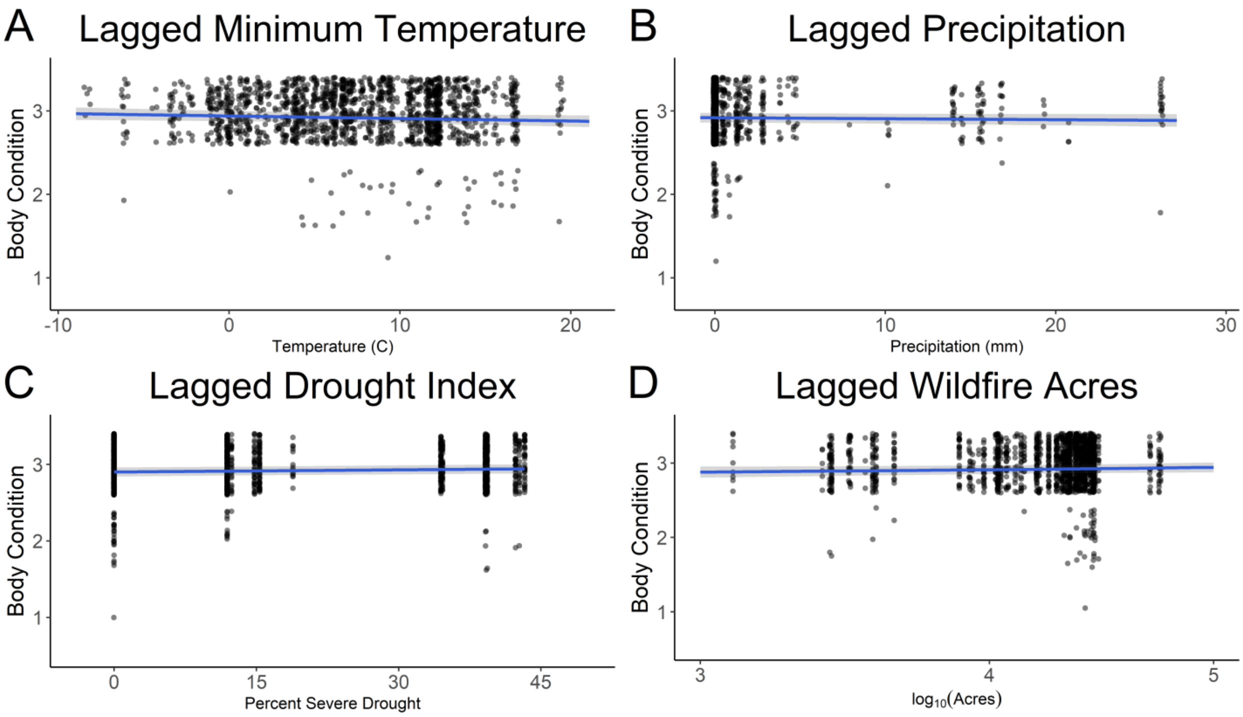


[1] While the statistical results noted in the paper for Body Condition come from a PGLMM, in order to be able to visualize these trends, we used a LMM instead, which did not produce qualitatively different results.

**FIGURE S4.** Trends in the analysis of covariance between $\delta^2$H isotope values and **A**) Julian day and species, **B**) Year and species, and **C**) the interaction between Julian day and year based on feathers collected from four species of birds at the Rio Mesa banding station in Utah (2021-2023). GWCS = Gambel's White-crowned Sparrow, WETA = Western Tanager, WIWA = Wilson's Warbler, and WWVI = Western Warbling-Vireo.

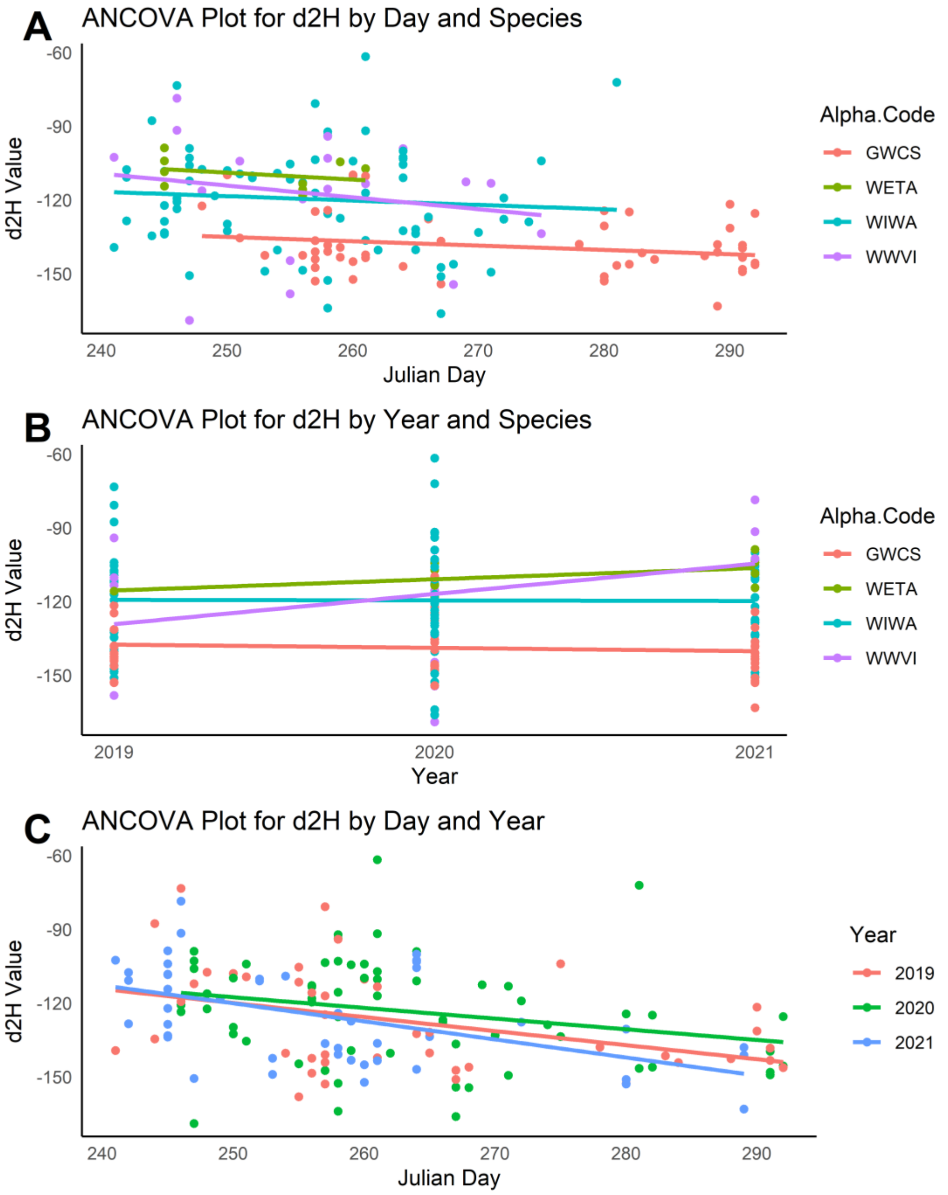

**FIGURE S5.** Non-metric Multidimensional Scaling (NMDS) plots between the 90% weighted distance estimates ("w90Dist"), Julian day, and year for four species of birds: **A**) WWVI = Western Warbling-Vireo, **B**) GWCS = Gambel's White-crowned Sparrow, **C**) WIWA = Wilson's Warbler, and **D**) WETA = Western Tanager. For species with less than four points for a given year, convex hull polygons rather than NMDS ellipses are depicted. See Table S6 for results from analyses of variance between these variables for each species.

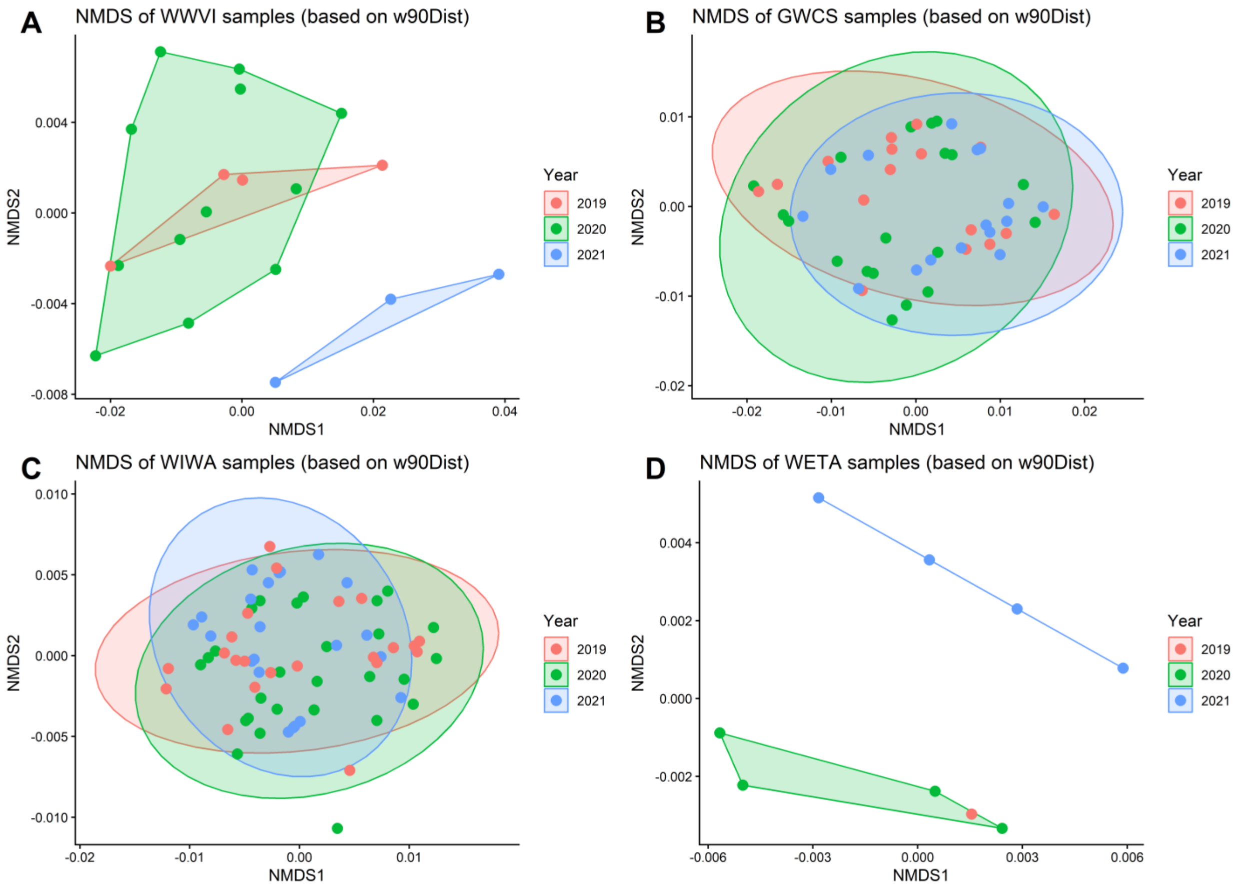

**FIGURE S6A.** Probability density plots of Western Warbling Vireos (*Vireo swainsoni*) captured during fall migration at the Rio Mesa bird banding station in southern Utah. Plots are generated based on stable hydrogen isotope analyses of feathers collected from the birds. The plots depicted here represent birds captured in 2020. Numbers correspond with individual sample IDs.

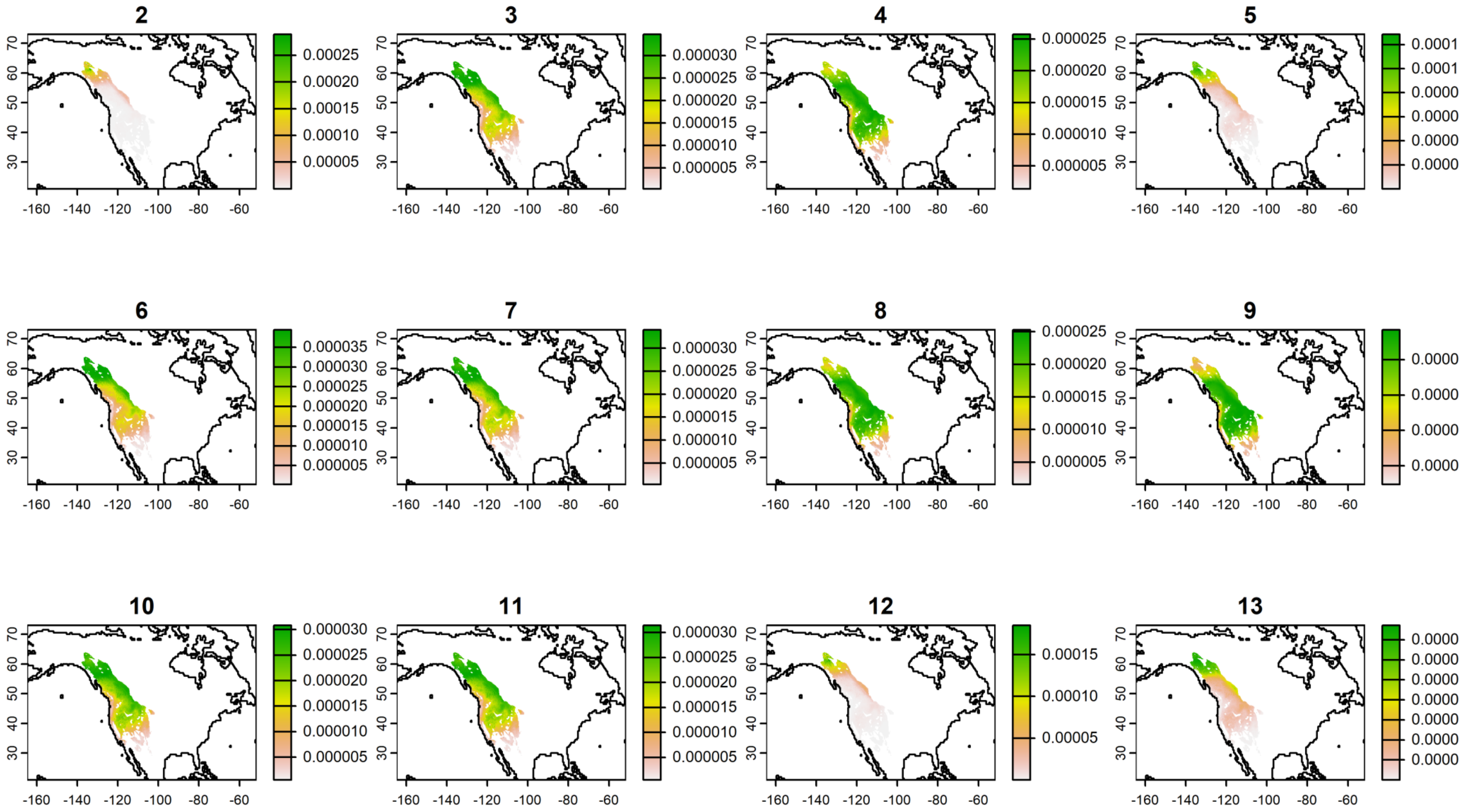

**FIGURE S6B.** Probability density plots of Western Warbling Vireos (*Vireo swainsoni*) captured during fall migration at the Rio Mesa bird banding station in southern Utah. Plots are generated based on stable hydrogen isotope analyses of feathers collected from the birds. Plots on the top row represent birds captured in 2021, while the other four plots are of birds captured in 2019. Numbers correspond with individual sample IDs.

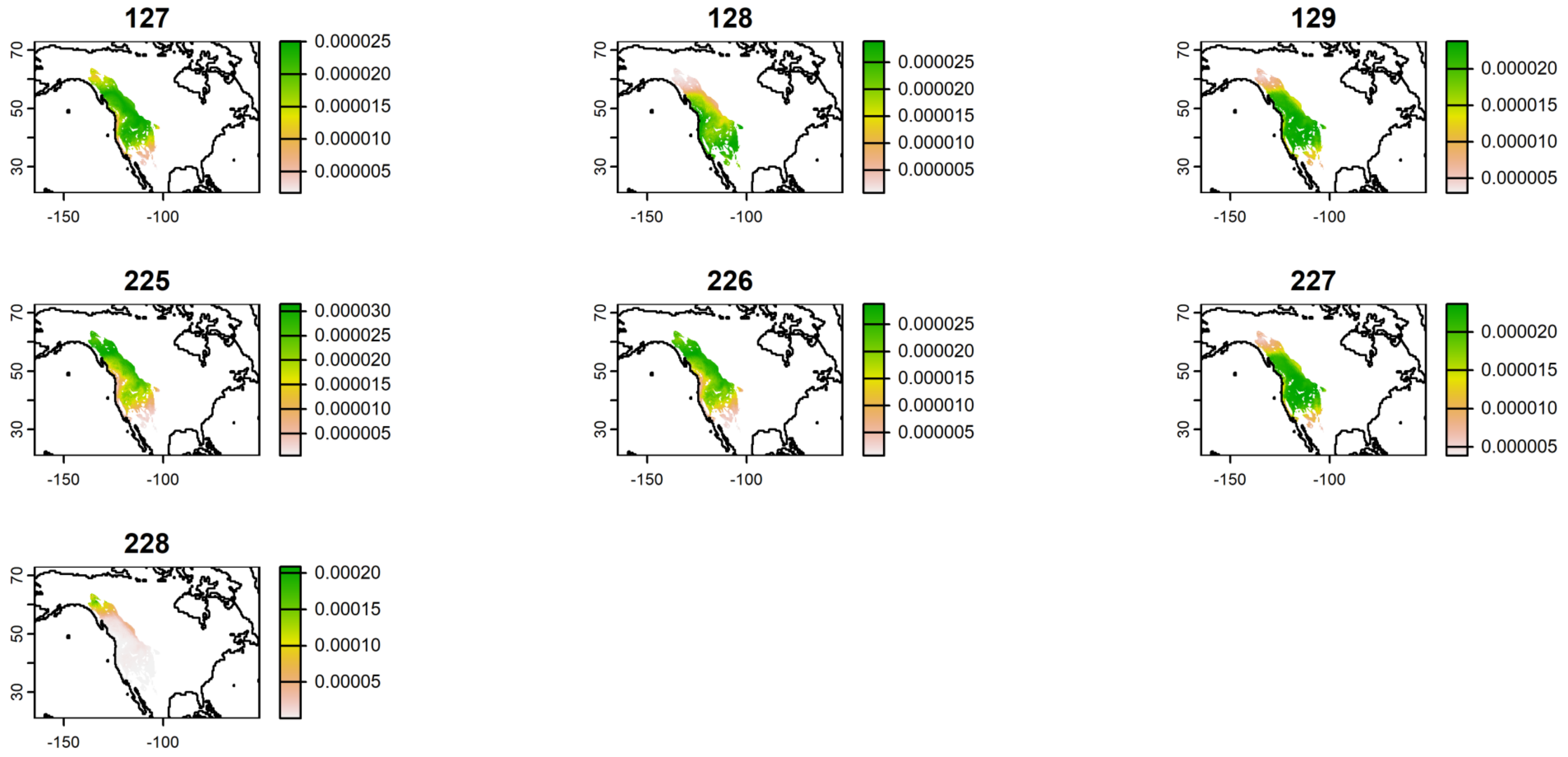

**FIGURE S6A.** Probability density plots of Gambel's White-crowned Sparrows (*Zonotrichia leucocphrys gambelii*) captured during fall migration at the Rio Mesa bird banding station in southern Utah. Plots are generated based on stable hydrogen isotope analyses of feathers collected from the birds. The plots depicted here represent birds captured in 2019. Numbers correspond with individual sample IDs.

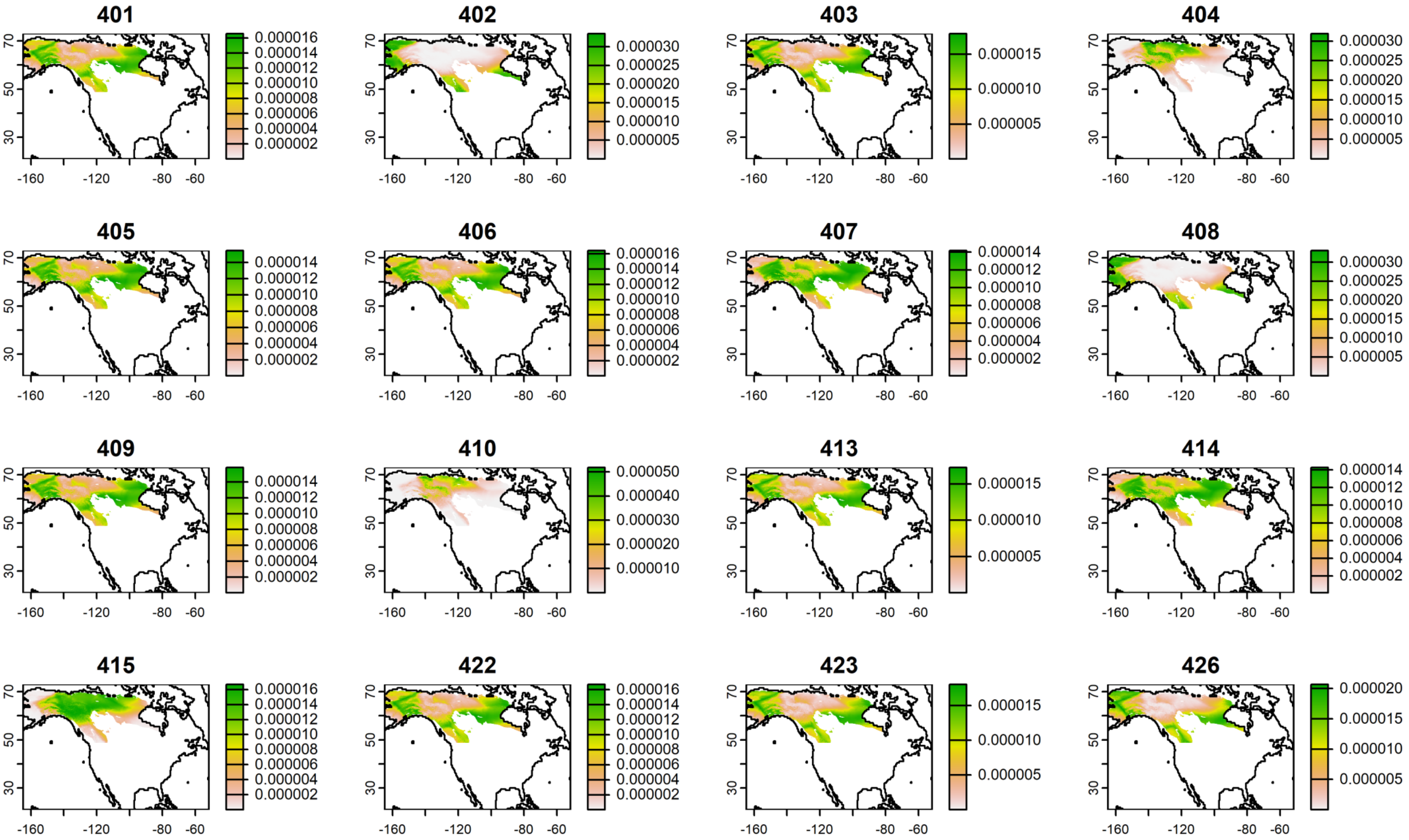

**FIGURE S6B.** Probability density plots of Gambel's White-crowned Sparrows (*Zonotrichia leucocphrys gambelii*) captured during fall migration at the Rio Mesa bird banding station in southern Utah. The plots depicted here represent birds captured in 2020. Numbers correspond with individual sample IDs. Birds in plots 35, 433, and 434 appear to have originated from the region consisting of Wrangell-St. Elias National Park in Alaska and Kluane National Park Reserve in the Yukon.

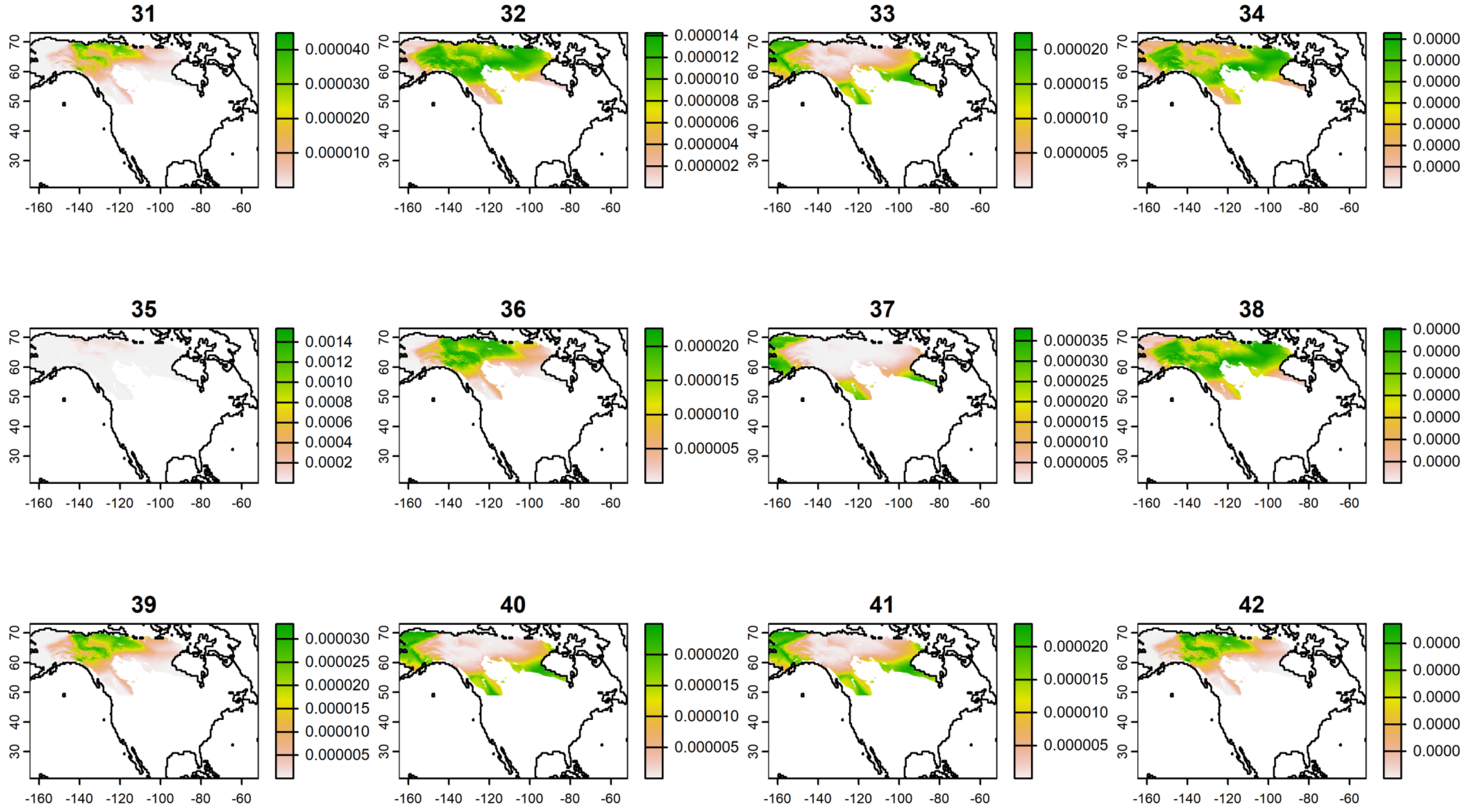

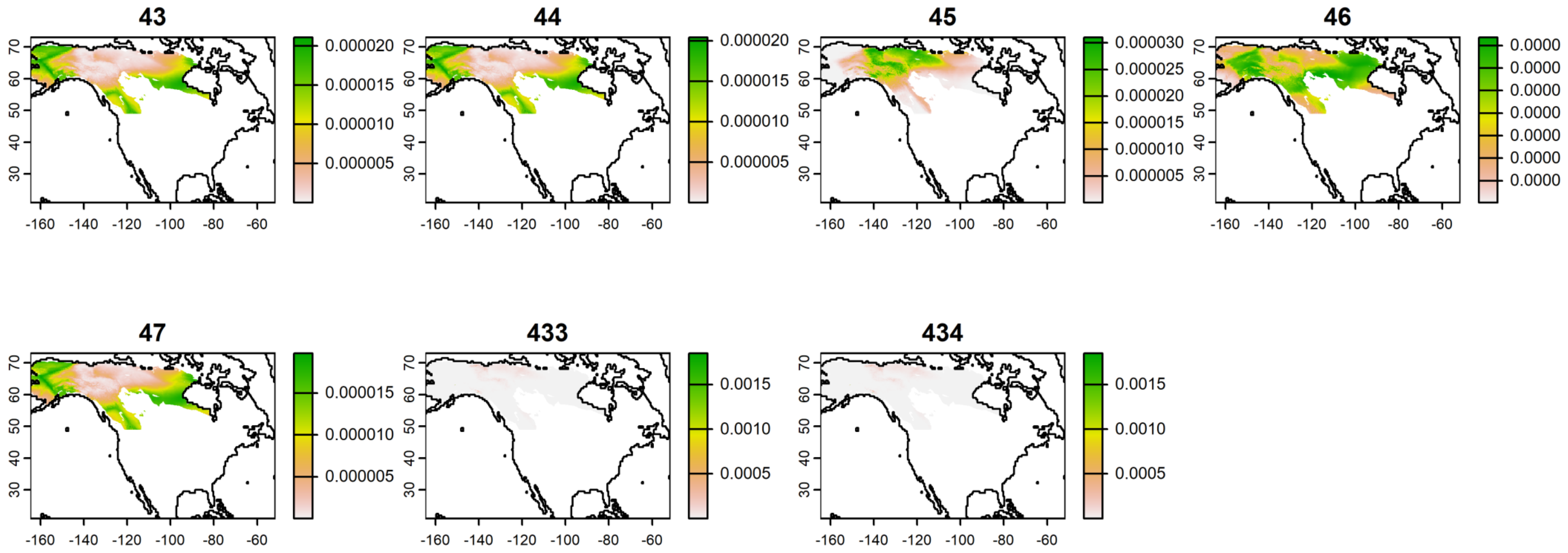
43
44
45
46
47
433
434
70
60
50
40
30
-160 -140 -120 -100 -80 -60
0.000020
0.000015
0.000010
0.000005
0.000030
0.000025
0.000020
0.000015
0.000010
0.000005
0.0000
0.0015
0.0010
0.0005

**FIGURE S6C.** Probability density plots of Gambel's White-crowned Sparrows (*Zonotrichia leucocphrys gambelii*) captured during fall migration at the Rio Mesa bird banding station in southern Utah. The plots depicted here represent birds captured in 2021. Numbers correspond with individual sample IDs.

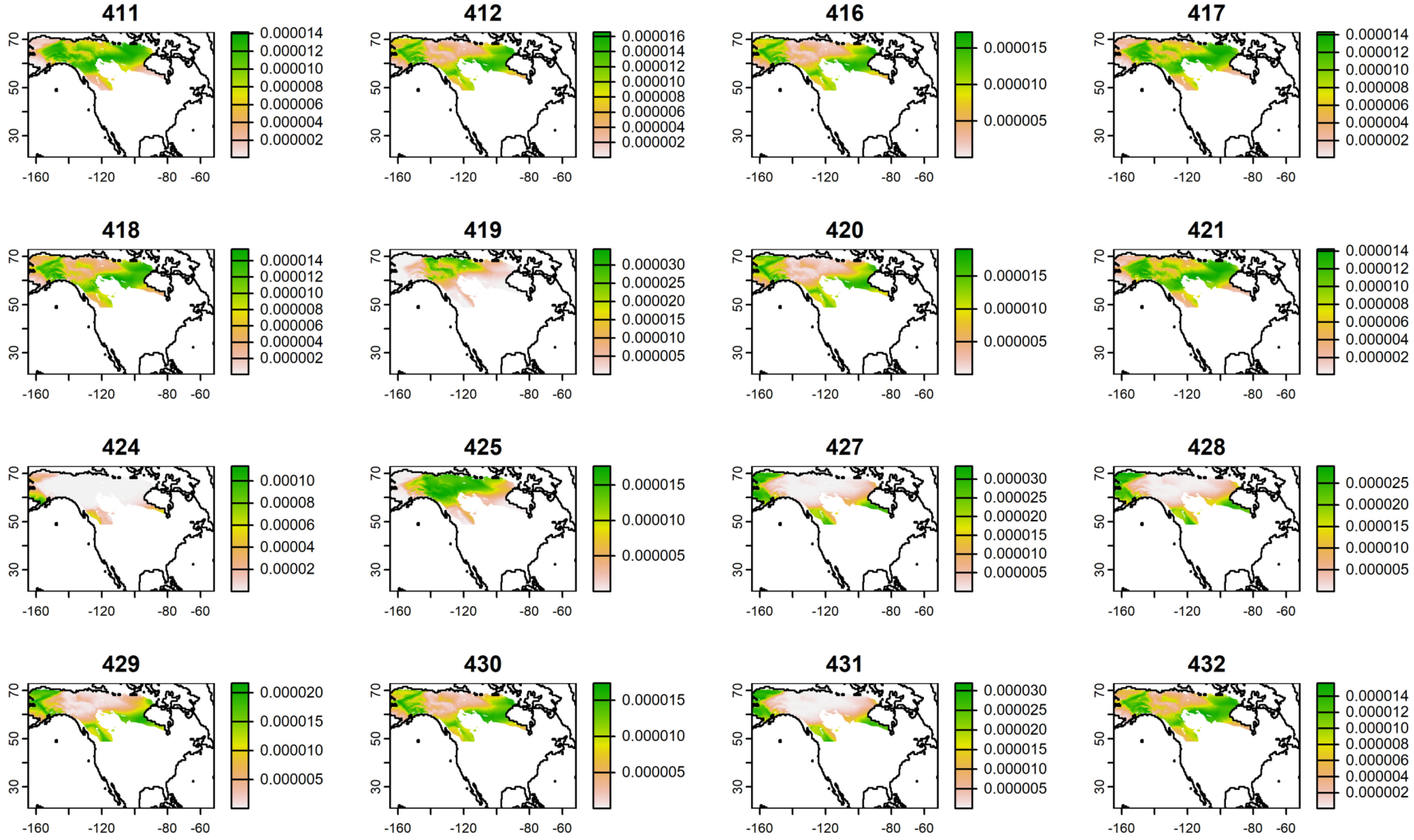

**FIGURE S7A.** Probability density plots of Wilson's Warblers (*Cardellina pusilla*) captured during fall migration at the Rio Mesa bird banding station in southern Utah. The plots depicted here represent birds captured in 2019. Numbers correspond with individual sample IDs.

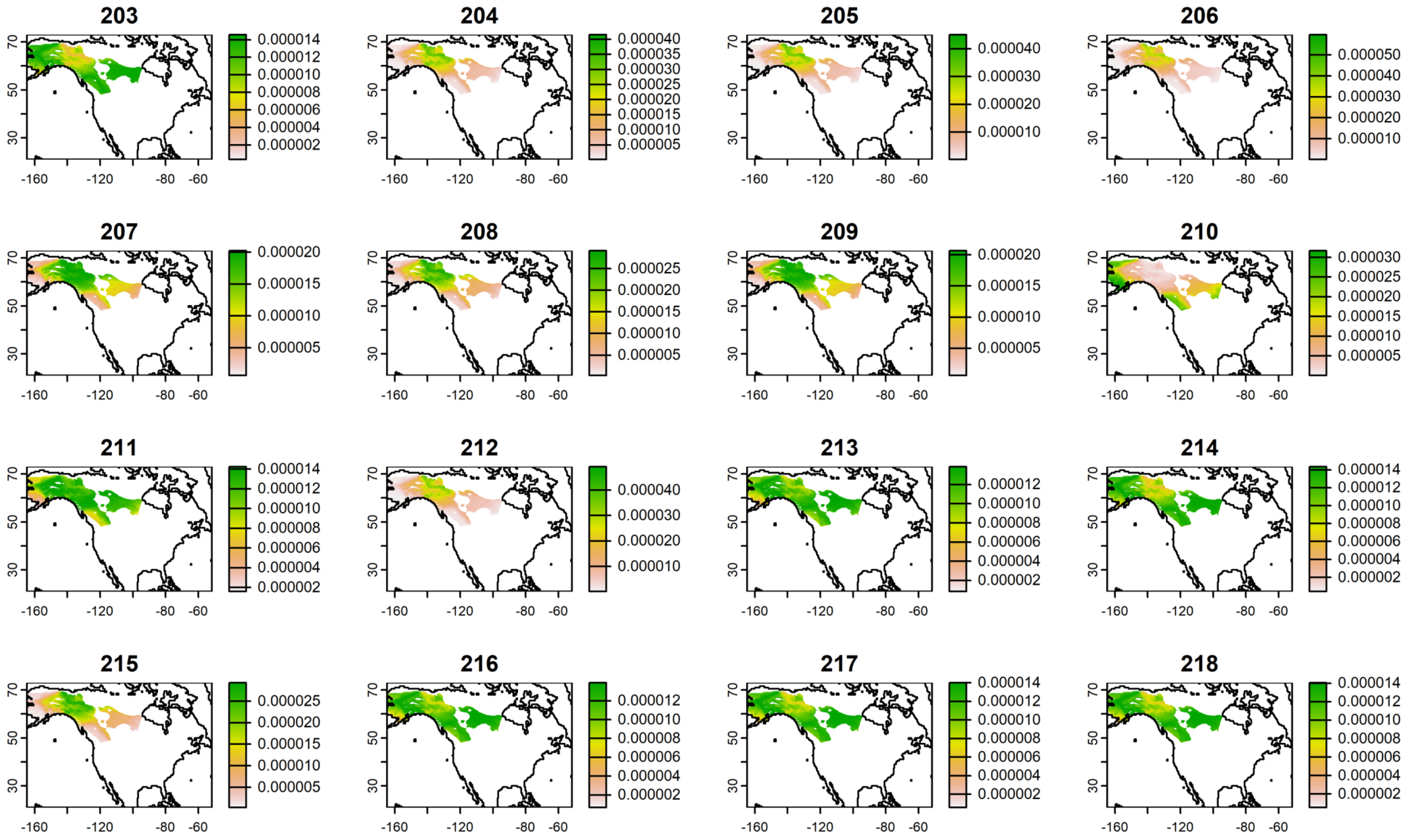

**219**

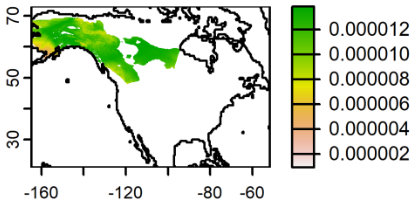


**220**

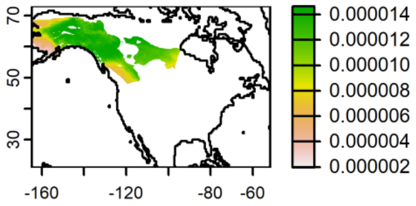


**221**

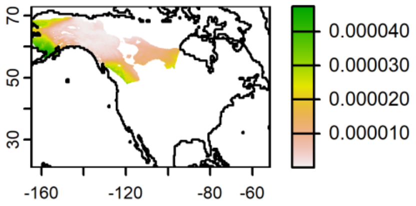


**222**

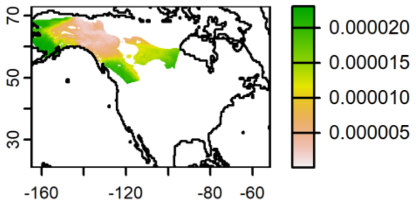


**223**

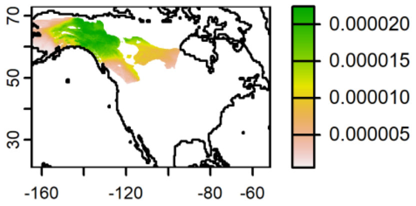


**224**

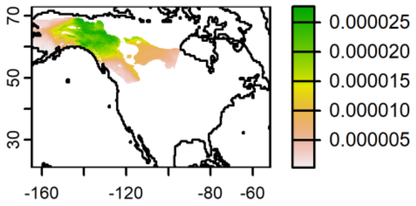

**FIGURE S7B.** Probability density plots of Wilson's Warblers (*Cardellina pusilla*) captured during fall migration at the Rio Mesa bird banding station in southern Utah. The plots depicted here represent birds captured in 2020. Numbers correspond with individual sample IDs. Birds 21 and 58 appear to have originated from the region consisting of Wrangell-St. Elias National Park in Alaska and Kluane National Park Reserve in the Yukon.

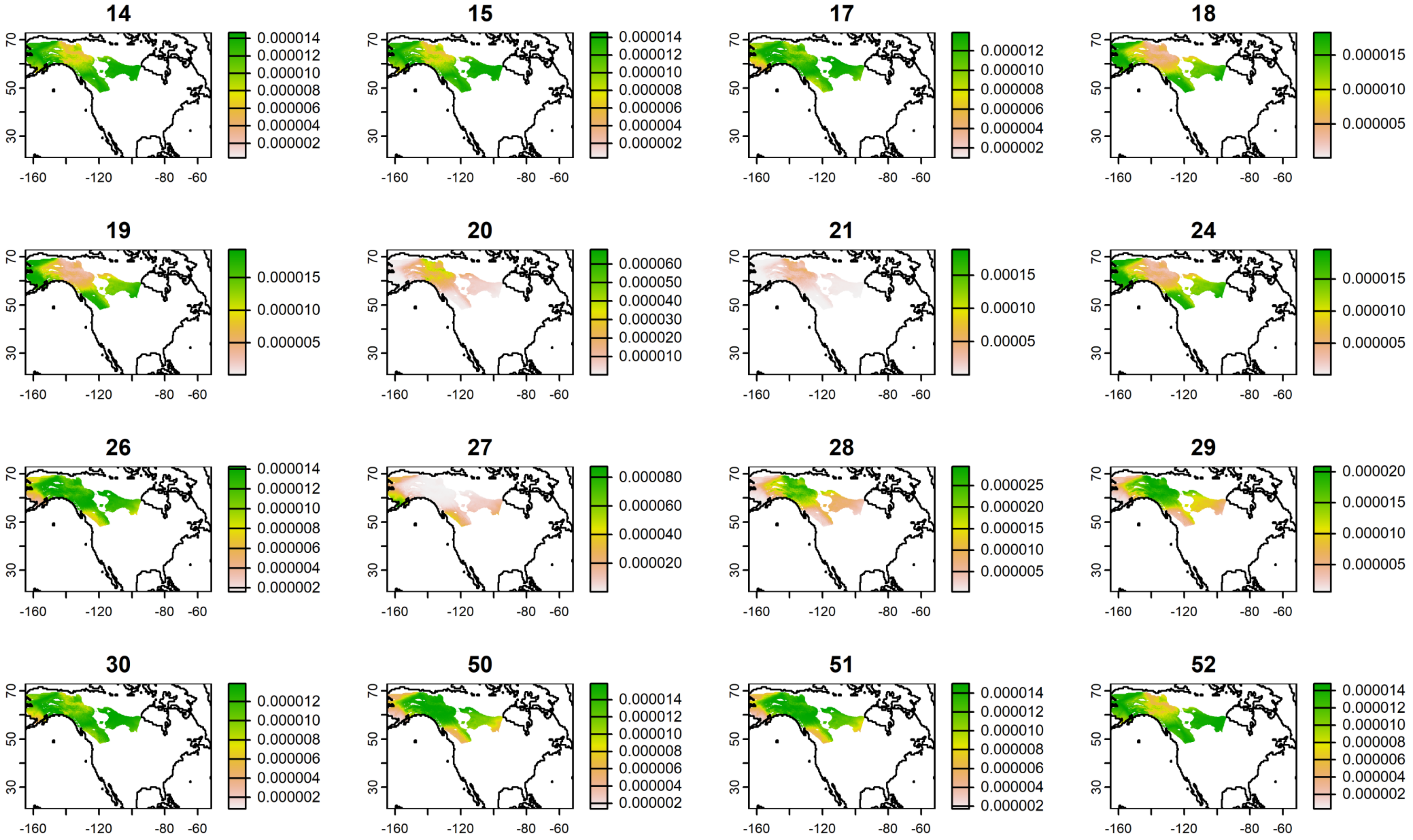

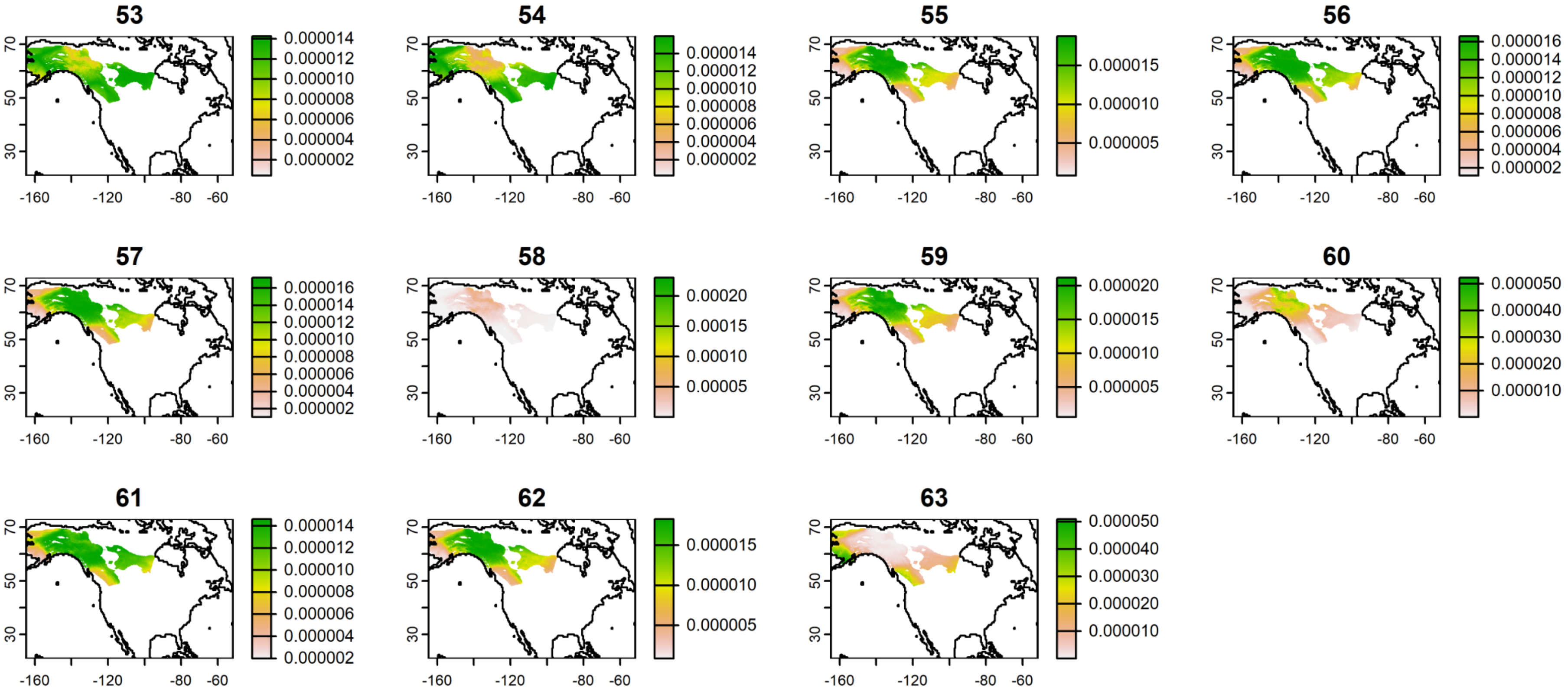
53
0.000014
0.000012
0.000010
0.000008
0.000006
0.000004
0.000002
54
0.000014
0.000012
0.000010
0.000008
0.000006
0.000004
0.000002
55
0.000015
0.000010
0.000005
56
0.000016
0.000014
0.000012
0.000010
0.000008
0.000006
0.000004
0.000002
57
0.000016
0.000014
0.000012
0.000010
0.000008
0.000006
0.000004
0.000002
58
0.00020
0.00015
0.00010
0.00005
59
0.000020
0.000015
0.000010
0.000005
60
0.000050
0.000040
0.000030
0.000020
0.000010
61
0.000014
0.000012
0.000010
0.000008
0.000006
0.000004
0.000002
62
0.000015
0.000010
0.000005
63
0.000050
0.000040
0.000030
0.000020
0.000010
70
50
30
-160
-120
-80
-60

**FIGURE S7C.** Probability density plots of Wilson's Warblers (*Cardellina pusilla*) captured during fall migration at the Rio Mesa bird banding station in southern Utah. The plots depicted here represent birds captured in 2021. Numbers correspond with individual sample IDs.

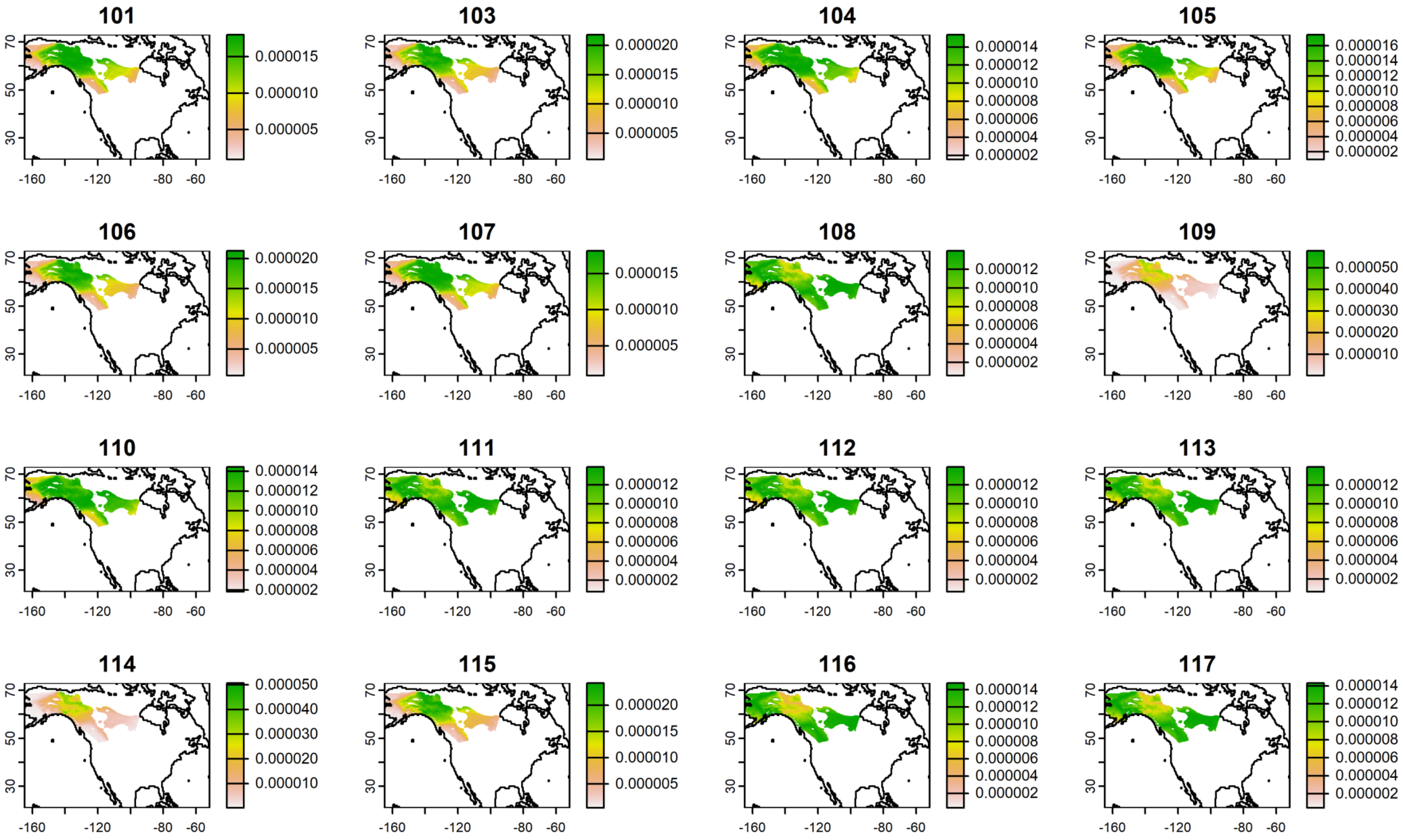

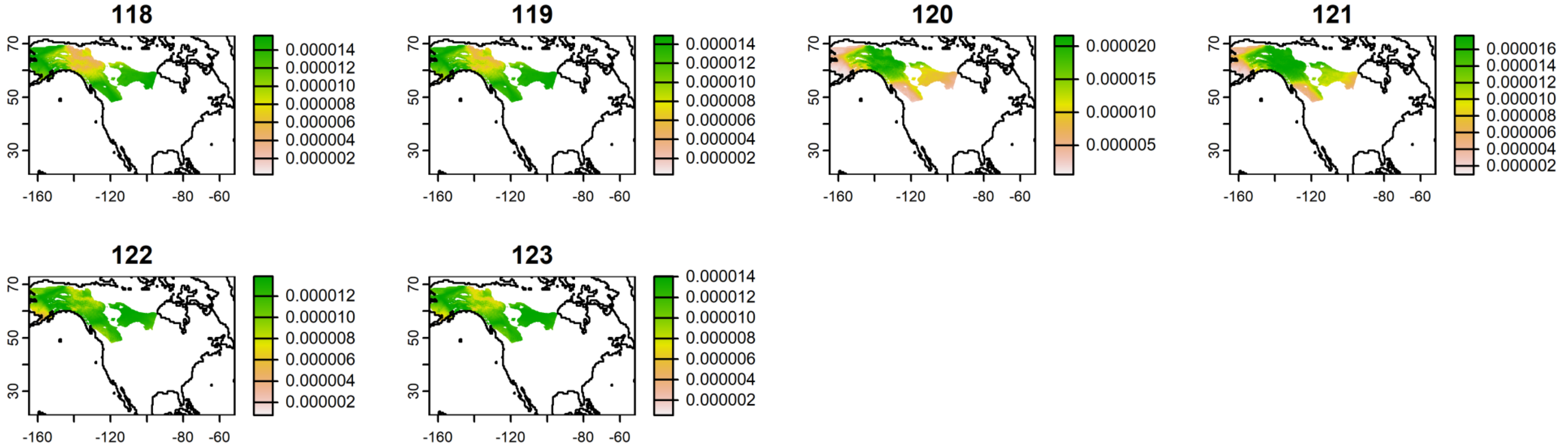
118
0.000014
0.000012
0.000010
0.000008
0.000006
0.000004
0.000002
119
0.000014
0.000012
0.000010
0.000008
0.000006
0.000004
0.000002
120
0.000020
0.000015
0.000010
0.000005
121
0.000016
0.000014
0.000012
0.000010
0.000008
0.000006
0.000004
0.000002
122
0.000012
0.000010
0.000008
0.000006
0.000004
0.000002
123
0.000014
0.000012
0.000010
0.000008
0.000006
0.000004
0.000002
70
50
30
-160
-120
-80
-60

**FIGURE S8.** Probability density plots of Western Tanagers (*Piranga ludoviciana*) captured during fall migration at the Rio Mesa bird banding station in southern Utah. The plots on the top row represent birds captured in 2020, the second row for birds captured in 2021, and the lone plot on the third row is of a bird captured in 2019. Numbers correspond with individual sample IDs.

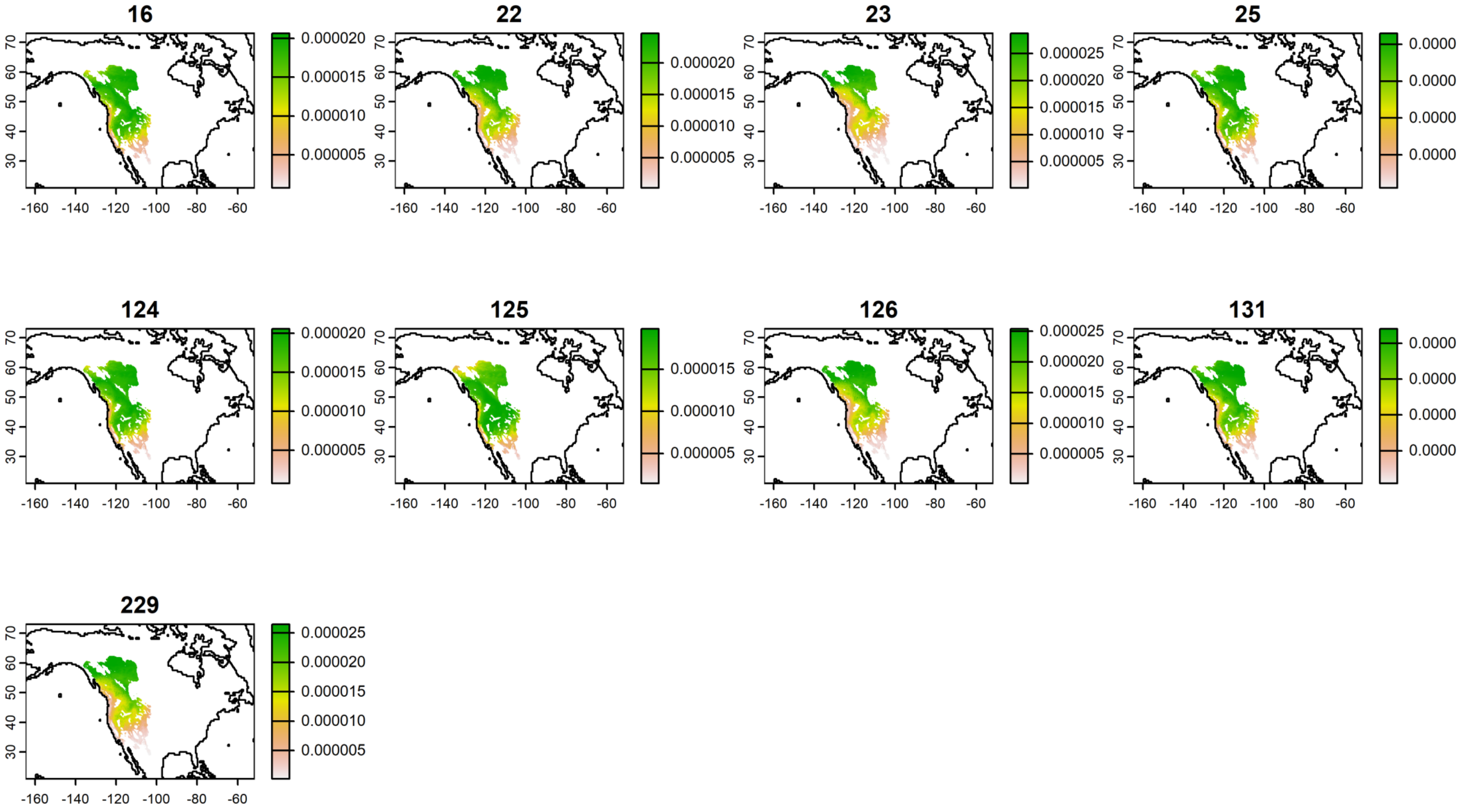

**TABLE S1**. Temporal trends across our study period (2019-2023) for our environmental variables. Results come from generalized linear models, where the intercept represents the mean value of each environmental variable; SE = standard error, T_min = minimum temperature, Precip = precipitation, and Fire = daily estimate of wildfire acres. See the Methods for information on the sources of and specifications for these various variables. Visualizations of these temporal trends for climate variables at both locations can be seen in Figure S2.

| Variable | Interaction Term | Estimate | SE | t value | *p* value |
|---|---|---|---|---|---|
| T_min | Intercept | 4.267 | 0.767 | 5.567 | <0.001 |
| | Year | 0.273 | 0.229 | 1.191 | 0.234 |
| Precip | Intercept | 0.580 | 0.501 | 1.158 | 0.248 |
| | Year | 0.162 | 0.150 | 1.082 | 0.280 |
| Drought | Intercept | 11.465 | 2.102 | 5.454 | <0.001 |
| | Year | 1.123 | 0.628 | 1.786 | 0.075 |
| Fire | Intercept | 3.967 | 0.080 | 49.572 | <0.001 |
| | Year | 0.050 | 0.024 | 2.085 | 0.038 |

**TABLE S2**. The effects of environmental and temporal variables on the daily number of birds captured at Rio Mesa, Utah (n= 3396) across a five-year period (2019-2023). Results come from a linear mixed-effects model, where the intercept represents the mean value of each fixed effect variable. SE = standard error, DF = degrees of freedom, T_min = minimum temperature, Precip = precipitation, Fire = daily estimate of wildfire acres, Julian = Julian day, and Julian:Year is the interaction between these two variables.

| Term | Estimate | SE | DF | t value | p value |
|---|---|---|---|---|---|
| Intercept | -6.989 | 3.300 | 252.513 | -2.118 | 0.035 |
| T_min | 0.793 | 0.722 | 253.454 | 1.099 | 0.273 |
| Precip | 0.109 | 0.403 | 250.384 | 0.272 | 0.786 |
| Drought | 0.869 | 0.470 | 254.941 | 1.849 | 0.066 |
| Fire | 4.712 | 0.794 | 254.451 | 5.934 | <0.001 |
| Julian | 0.033 | 0.040 | 253.861 | 0.823 | 0.411 |
| Year | -2.016 | 0.435 | 249.837 | -4.640 | <0.001 |
| Julian:Year | 0.041 | 0.023 | 254.818 | 1.744 | 0.082 |

**TABLE S3**. The effects of environmental and temporal variables on the body masses of birds captured at Rio Mesa, Utah (n= 3094) across a five-year period (2019-2023). Results come from a phylogenetic generalized linear mixed effects model (PGLMM), where the intercept represents the mean value of each fixed effect variable. SE = standard error, T_min_lag = lagged minimum temperature, Precip_lag = lagged precipitation, Drought_lag = lagged drought index, Fire_lag = lagged daily estimate of wildfire acres, Julian = Julian day, and Julian:Year is the interaction between these two variables. The lagged environmental variables represent values one week prior to an individual bird's capture.

| Term | Estimate | SE | Z score | p value |
|---|---|---|---|---|
| Intercept | 0.961 | 0.020 | 47.284 | <0.001 |
| T_min_lag | 0.001 | 0.002 | 0.273 | 0.785 |
| Precip_lag | <0.001 | 0.001 | 0.139 | 0.889 |
| Drought_lag | -0.001 | 0.001 | -0.597 | 0.551 |
| Fire_lag | -0.016 | 0.002 | -6.692 | <0.001 |
| Julian | <0.001 | 0.000 | 3.108 | 0.002 |
| Year | -0.005 | 0.002 | -2.342 | 0.019 |
| Julian:Year | <0.001 | 0.000 | 0.069 | 0.945 |
| Fat Score | 0.023 | 0.001 | 21.589 | <0.001 |

**TABLE S4**. The effects of environmental and temporal variables on the body condition of birds captured at Rio Mesa, Utah (n= 1532) across a three-year period (2021-2023). Results come from a phylogenetic generalized linear mixed effects model (PGLMM), where the intercept represents the mean value of each fixed effect variable. SE = standard error, T_min_lag = lagged minimum temperature, Precip_lag = lagged precipitation, Drought_lag = lagged drought index, Fire_lag = lagged daily estimate of wildfire acres, Julian = Julian day, and Julian:Year is the interaction between these two variables. The lagged environmental variables represent values one week prior to an individual bird's capture.

| **Term** | **Estimate** | **SE** | **Z score** | **p value** |
|---|---|---|---|---|
| Intercept | 2.733 | 0.117 | 23.375 | <0.001 |
| T_min_lag | -0.016 | 0.008 | -1.975 | 0.048 |
| Precip_lag | -0.005 | 0.004 | -1.221 | 0.222 |
| Drought_lag | 0.015 | 0.006 | 2.453 | 0.014 |
| Fire_lag | 0.032 | 0.020 | 1.607 | 0.108 |
| Julian | \|<0.001\|[2] | <0.001 | -0.688 | 0.492 |
| Year | 0.011 | 0.022 | 0.519 | 0.604 |
| Julian:Year | <0.001 | <0.001 | 0.733 | 0.464 |

[2] The absolute value here refers to a number that is greater than the -0.001 but less than zero.

**TABLE S5**. Results from analyses of variance (AOV) between $\delta^2$H isotope values and the interaction between year and Julian day for all four species separately and “all” together; the latter also included species itself as a covariate. WWVI = Western Warbling-Vireo, GWCS = Gambel’s White-crowned Sparrow, WIWA = Wilson’s Warbler, WETA = Western Tanager, and DF = degrees of freedom.

| Species | Term | DF | Sum of Squares | Mean Square | F value | p value |
|---|---|---|---|---|---|---|
| All | Year | 1 | 94 | 94 | 0.276 | 0.600 |
| | Julian | 1 | 8308 | 8308 | 24.326 | <0.001 |
| | Species | 3 | 7525 | 7525 | 7.345 | <0.001 |
| | Year:Julian | 1 | 11 | 11 | 0.033 | 0.857 |
| WWVI | Year | 1 | 1055 | 1055.1 | 2.128 | 0.165 |
| | Julian | 1 | 38 | 37.6 | 0.076 | 0.787 |
| | Year:Julian | 1 | 1944 | 1943.8 | 3.921 | 0.066 |
| GWCS | Year | 1 | 58 | 57.5 | 0.427 | 0.517 |
| | Julian | 1 | 415 | 414.6 | 3.076 | 0.086 |
| | Year:Julian | 1 | 65 | 65.4 | 0.485 | 0.490 |
| WIWA | Year | 1 | 2 | 2.1 | 0.005 | 0.946 |
| | Julian | 1 | 211 | 211.2 | 0.465 | 0.498 |
| | Year:Julian | 1 | 1345 | 1344.9 | 2.962 | 0.090 |
| WETA | Year | 1 | 83.72 | 83.72 | 2.475 | 0.176 |
| | Julian | 1 | 14.32 | 14.32 | 0.423 | 0.544 |
| | Year:Julian | 1 | 62.31 | 62.31 | 1.842 | 0.233 |

**TABLE S6**. Results from analyses of variance (AOV) between the weighted 90% distance estimates and the interaction between year and Julian day for all four species separately. WWVI = Western Warbling-Vireo, GWCS = Gambel's White-crowned Sparrow, WIWA = Wilson's Warbler, WETA = Western Tanager, and DF = degrees of freedom.

| Species | Term | DF | Sum of Squares | Mean Square | F value | p value |
|---|---|---|---|---|---|---|
| WWVI | Year | 1 | 305142 | 305142 | 3.758 | 0.072 |
| | Julian | 1 | 86332 | 86332 | 1.063 | 0.319 |
| | Year:Julian | 1 | 335557 | 335557 | 4.132 | 0.060 |
| GWCS | Year | 1 | 15507 | 15507 | 0.223 | 0.639 |
| | Julian | 1 | 87234 | 87234 | 1.254 | 0.268 |
| | Year:Julian | 1 | 20605 | 20605 | 0.297 | 0.589 |
| WIWA | Year | 1 | 13 | 13 | 0.001 | 0.981 |
| | Julian | 1 | 13834 | 13834 | 0.630 | 0.430 |
| | Year:Julian | 1 | 65419 | 65419 | 2.980 | 0.089 |
| WETA | Year | 1 | 9817 | 9817 | 2.328 | 0.188 |
| | Julian | 1 | 1623 | 1623 | 0.385 | 0.562 |
| | Year:Julian | 1 | 6820 | 6820 | 1.617 | 0.259 |

**TABLE S7**. The effects of environmental and temporal variables on the body masses of birds captured at Rio Mesa, Utah (n= 3094) across a five-year period (2019-2023). Results come from a linear mixed-effects model, where the intercept represents the mean value of each fixed effect variable. SE = standard error, DF = degrees of freedom, T_min_lag = lagged minimum temperature, Precip_lag = lagged precipitation, Drought_lag = lagged drought index, Fire_lag = lagged daily estimate of wildfire acres, Julian = Julian day, and Julian:Year is the interaction between these two variables. The lagged environmental variables represent values one week prior to an individual bird's capture.

| **Term** | **Estimate** | **SE** | **DF** | **t value** | **p value** |
|---|---:|---:|---:|---:|---:|
| Intercept | 0.950 | 0.018 | 26.710 | 54.035 | <0.001 |
| T_min_lag | 0.001 | 0.002 | 3017.000 | 0.289 | 0.773 |
| Precip_lag | <0.001 | 0.001 | 3012.000 | 0.123 | 0.902 |
| Drought_lag | -0.001 | 0.001 | 2894.000 | -0.678 | 0.498 |
| Fire_lag | -0.016 | 0.002 | 2800.000 | -6.661 | 0.000 |
| Julian | <0.001 | <0.001 | 3045.000 | 3.165 | 0.002 |
| Year | -0.007 | 0.003 | 10.580 | -2.520 | 0.029 |
| Julian:Year | $\lvert<0.001\rvert$[3] | <0.001 | 122.400 | -0.051 | 0.959 |
| Fat Score | 0.023 | 0.001 | 3024.000 | 21.582 | <0.001 |

[3] The absolute value here refers to a number that is greater than the -0.001 but less than zero.

**TABLE S8**. The effects of environmental and temporal variables on the body condition of birds captured at Rio Mesa, Utah (n= 1532) across a three-year period (2021-2023). Results come from a linear mixed-effects model, where the intercept represents the mean value of each fixed effect variable. SE = standard error, DF = degrees of freedom, T_min_lag = lagged minimum temperature, Precip_lag = lagged precipitation, Drought_lag = lagged drought index, Fire_lag = lagged daily estimate of wildfire acres, Julian = Julian day, and Julian:Year is the interaction between these two variables. The lagged environmental variables represent values one week prior to an individual bird's capture.

| Term | Estimate | SE | DF | t value | p value |
|---|---|---|---|---|---|
| Intercept | 2.784 | 0.088 | 631.300 | 31.560 | <0.001 |
| T_min_lag | -0.016 | 0.008 | 1375.000 | -2.005 | 0.045 |
| Precip_lag | -0.005 | 0.004 | 1410.000 | -1.241 | 0.215 |
| Drought_lag | 0.016 | 0.006 | 1443.000 | 2.517 | 0.012 |
| Fire_lag | 0.032 | 0.020 | 1348.000 | 1.608 | 0.108 |
| Julian | \|<0.001\|[4] | <0.001 | 1476.000 | -0.645 | 0.519 |
| Year | 0.004 | 0.023 | 28.030 | 0.159 | 0.875 |
| Julian:Year | <0.001 | <0.001 | 1399.000 | 0.642 | 0.521 |

[4] The absolute value here refers to a number that is greater than the -0.001 but less than zero.
